# The south Congo Basin was critical to Bantu settlement of south central Africa

**DOI:** 10.64898/2026.08.14.741591

**Authors:** Jeremy Choin, Kathryn de Luna, Jeffrey Fleisher, Elizabeth Sawchuk, Steven Goldstein, Potiphar Kaliba, Maggie Katongo, Alan G. Morris, George Mudenda, Menno Welling, Kendra Sirak, Kim Callan, Lora Iliev, Ann Marie Lawson, Megan Michel, Jonas Oppenheimer, Lijun Qiu, J. Noah Workman, Aisling Kearn, Matthew Mah, Gregory Soos, Swapan Mallick, Nadin Rohland, Jessica Thompson, Mary E. Prendergast, David Reich

**Author notes:** Corresponding authors: Jeremy Choin, Mary Prendergast and David Reich.

## Abstract

South central Africa, between the Congo Basin, the Great Lakes, and southern Africa, has long served as a corridor for human movement. Yet, it remains unclear whether there is genetic continuity between pre-Iron Age foragers and later Bantu-associated populations and whether the settlement of this region reflects multiple Bantu-associated migrations rather than the simpler serial founder model suggested by existing genomic data. To do so, we generated genome-wide ancient DNA from 71 Iron Age and historical individuals from present-day Zambia and Malawi, and analyzed these genomes alongside published data from present-day Africans. One late Iron Age individual (16^th^-17^th^ century) from Kalala Island in the Kafue River, Zambia, carries 40% non Bantu related ancestry that closely matches local Later Stone Age foragers. Admixture for this individual is estimated to be 850 years ago, several centuries earlier than reported for present day BaTwa from the same region. Focusing on the Bantu-related ancestry, haplotype-based analyses identify two main clusters among Iron Age, historical, and present-day south central Bantu groups associated with different Bantu-related migrations. These results reveal a layered history in which at least two Bantu expansions radiated from the southern Congo Basin, with south central Africa acting both as a crossroads of these movements and as a staging area for the subsequent southward expansion toward southern Africa.

## Introduction

South central Africa, encompassing much of present-day Malawi and Zambia, occupies a key geographic position between the Congo Basin, the Great Lakes region, and the southern African deserts. This area has long functioned as a corridor for human mobility and exchange, contributing to repeated episodes of population movement and contact from the Pleistocene through the historical period. Archaeological and genomic evidence indicates that the region was inhabited by diverse hunter-gatherer populations throughout the Late Pleistocene and Holocene, associated with Later Stone Age (LSA) stone tool traditions^1–7^. From at least the early first millennium CE, farming and iron-working communities spread into present-day Malawi and Zambia, a process broadly linked to the expansion of Bantu language-speaking populations^8,9^ who spread out from western Africa beginning ∼4000 CE, albeit with important regional caveats^10^. There is wide agreement that this spread drastically changed the linguistic, cultural, and genetic landscapes. There is wide agreement that this spread drastically changed the linguistic, cultural, and genetic landscapes of much of Africa, but the precise mechanisms of expansion remain debated. Original assumptions of a unidirectional single expansion in the deeper past are now being challenged by models emphasizing multiple complex dispersals of Bantu speakers over millennia^8–11^. By the second half of the first millennium CE, transregional trade networks, shared currencies, and dispersed polities connected populations throughout and beyond the region, stretching across southern Africa and the Indian Ocean world. Archaeological evidence indicates that these processes involved continuous demographic movement and contact and mobility^12,13^; regional language data evince complex histories of contact, borrowing, layering, and absorption^8,9,14^. Late second millennium patterns of mobility, trade, and social organization were shaped by regional polities such as the Maravi state in Malawi, the powerful central African complexes associated with the Luba and Lunda traditions and healing cults and many others, as well as growing transcontinental trade (including in slaves)^15–19^.

Recent genomic studies, based almost exclusively on present-day people, have identified south central Africa as a pivotal region in the Bantu expansion. Previous work suggested that this expansion followed a series of founder events originating in western Africa south of the equatorial rainforest, with Bantu-speaking groups branching south toward Namibia and east through the regions surrounding Zambia into eastern and southern Africa^20,21^. More recently, Fortes-Lima et al.^22^, leveraging a more comprehensive genomic dataset, proposed that the Democratic Republic of Congo and Zambia formed an interaction zone, although barriers to gene flow within south central Africa left unresolved whether the region functioned primarily as a true crossroads of multiple Bantu migrations or as a splitting point along a serial founder route. In parallel, genetic studies of present-day and ancient African populations have shown that Bantu-speaking groups admixed to varying degrees with local communities encountered during their movement and settlement across the continent. In south central Africa, however, genomic studies of present-day populations generally indicate relatively low levels of detectable admixture with local forager groups. An important exception is provided by genetic studies of BaTwa-associated populations from the region^23^, which reveal evidence of admixture between Bantu-speaking groups and ancestry related to a now-extinct form of forager lineage, pointing to the persistence of forager-related genetic components in specific communities. Bioarchaeological analyses of LSA burials in Malawi and Zambia further suggest the presence of a central African forager ancestry unrelated to present-day groups^24–26^. While these lines of evidence indicate complex population history, studies of ancient DNA from this region have been rare, limiting interpretation of present-day genomes.

As a result, key questions remain unresolved: how much genetic continuity exists between pre-Iron Age foragers and later populations in Malawi and Zambia? Did the settlement of the region involve one or multiple waves of migration by genetically distinct Bantu-associated people? To address these questions, we generated genome-wide ancient DNA data from Iron Age and historical individuals from south central Africa and co-analyzed them with published ancient and present-day genomic data from across the continent. In addition, we investigated fine-scale population structure using previously published genome-wide data from present-day African populations, with a particular focus on Bantu-speaking groups and dense sampling from Malawi, Zambia, and the Democratic Republic of Congo (DRC)^22^.

### New Iron-age and Historical south central African DNA data

Numerous Iron Age sites across present-day Zambia and Malawi have human burials, but many of these are poorly preserved, fragmentary, and commingled in legacy collections lacking critical context information, dates, or bioarchaeological assessment (**Supplementary Material**). We began with archival research and an inventory of skeletal collections in the Livingstone Museum (Zambia) to select appropriate remains for analysis; we also sampled burials from both past and recent excavations in Malawi. In **Supplementary Material**, we provide research histories, contexts, previously published dates, and bioarchaeological assessments of these individuals.

We sampled 111 petrous bones, long bones, and teeth from 39 Iron Age and historic archaeological sites across both countries (**Supplementary Material, Supplementary Table 1**). In dedicated clean-room facilities, we generated new ancient DNA data from 76 of these samples, which represent 71 individuals. After quality control, we obtained analyzable data for 62 individuals (> 10,000 SNPs), with number of SNPs ranging from 10,546 to 514,173 (**Figure 1a and Supplementary Table 1**). We radiocarbon dated 55 samples from individuals producing readable ancient DNA via accelerator mass spectrometry (AMS). For many sites, this provided the first available dates or enabled a reassessment of published indirect, pre-AMS dates (**Supplementary Material**). Here we report calibrated Common Era (CE) dates using the SHCal20 curve^27^ in OxCal v4.4^28^, and report raw dates and isotopic values in **Supplementary Table 2**. We co-analyzed these newly sequenced and dated individuals with published ancient and present-day genome-wide data from Africa^22,29–39^. We also increased the coverage of three previously published Later Stone Age (LSA) individuals from Hora and Fingira sites^29^ (**Supplementary Table 1**)

**Figure 1.**
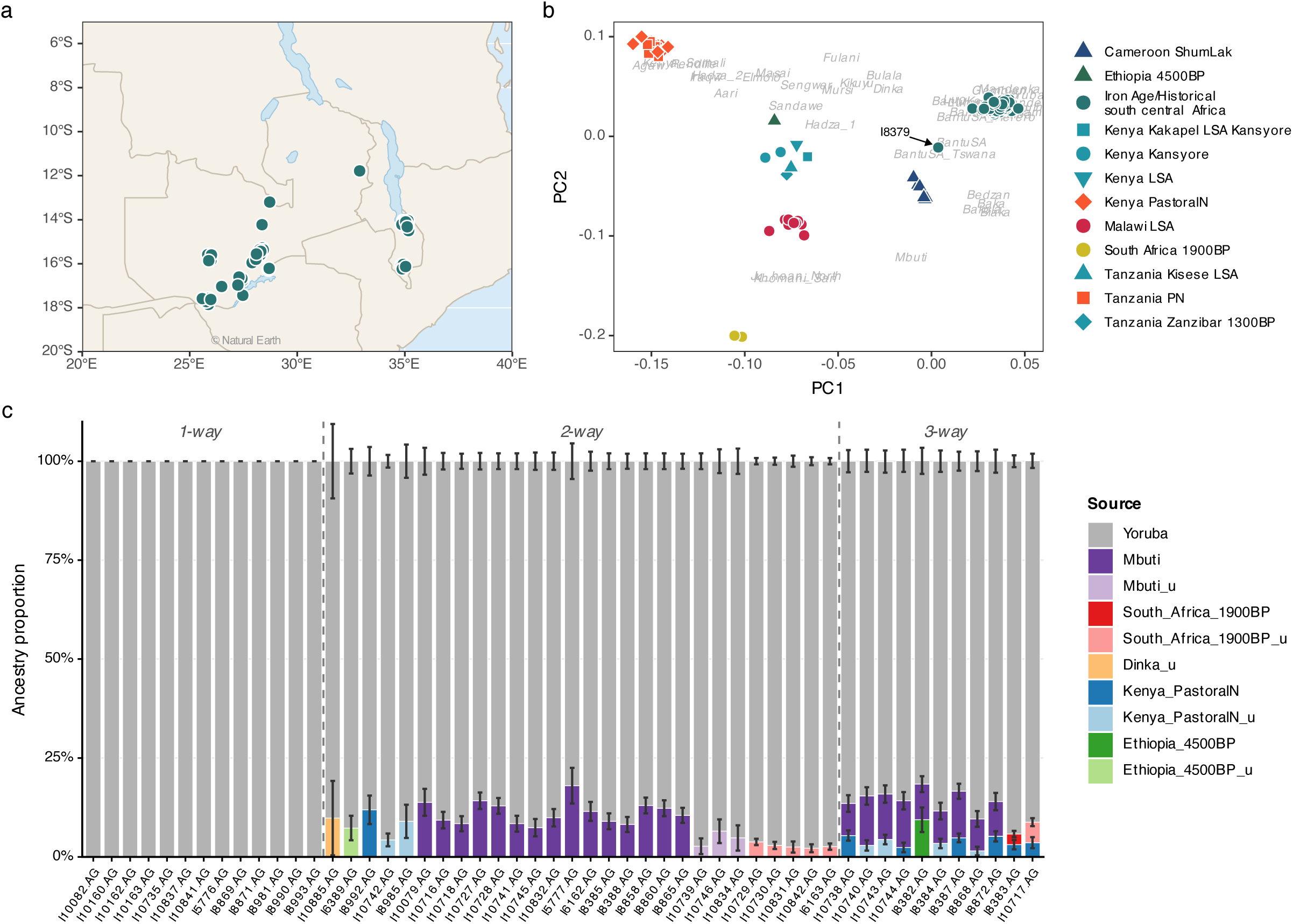
Ancient genomic diversity of Iron Age and Historical individuals from Malawi and Zambia. **(a)** Geographic locations of newly generated ancient individuals from Malawi and Zambia analyzed in this study. **(b)** Principal Component Analysis (PCA) computed on modern African populations (grey italic labels indicate population centroids) with ancient individuals projected onto the principal component axes (**Supplementary Table 3**). Ancient individuals are colored by group: Iron Age/historical south central Africa (teal), Malawi LSA (pink), eastern African foragers (teal-blue), pastoralists (orange), ancient Ethiopian hunter-gatherer (dark green), Cameroon hunter-gatherer (dark blue), and South African (yellow). The black arrow indicates the individual I8379 from Kalala Island (Kafue River, Zambia; 1516-1667 cal CE) **(c)** Distal ancestry modeling using qpAdm. Each bar represents the best-fitting admixture model for a given target individual under 1-way, 2-way, or 3-way models. Colors indicate ancestry sources as shown in the legend. Suffix “_u” denotes uncertain models, defined as either multiple passing models (p-value > 0.05) or a unique passing model with Z-score < 2 for the considered source; in the case of multiple passing models, the model with the lowest standard error is shown (**Supplementary Tables 4 and 5)**.

### Genetic affinity of ancient Malawians and Zambians with non-Bantu groups

Like linguistic and archaeological research, previous genomic studies of present-day sub-Saharan Africans have shown that Iron Age populations admixed to varying degrees with local African communities during their expansion, including: (i) central African rainforest populations, (ii) Nilo-Saharan and Afro-Asiatic pastoralists, (iii) eastern African foragers, and (iv) southern African foragers^20,23,33,40–43^. Although Bantu-related ancestry constitutes the dominant genetic component in south central Africa today^20,22,23^, the extent to which Iron Age populations from this region retain genetic contributions from non-Bantu-related groups remains unclear.

We first performed a principal component analysis (PCA) on the genomes of Iron Age and historical individuals from south central Africa generated in this study, together with published ancient and present-day genomes from Bantu- and non-Bantu-speaking populations from western, central (e.g., rainforest groups), eastern, and southern Africa (**Figure 1b and Supplementary Table 3**). Principal component 1 (PC1) captures genetic differentiation between Afro-Asiatic-speaking eastern Africans and southern African hunter-gatherers, while PC2 distinguishes West African from eastern African populations. Most Iron Age and historical individuals from south central Africa project toward the West African end of the PCA space, consistent with substantial Bantu-related ancestry. However, one individual, I8379 from Kalala Island (Kafue River, Zambia; 1516-1667 cal CE), an Iron Age archaeological context associated with fishing and hunting^5,44^, is notably shifted toward LSA south central African hunter-gatherers (**Figure 1b**).

To estimate ancestry proportions and likely distal sources, we applied qpWave and qpAdm^45–47^. We tested for the presence of ancestry related to eastern African foragers (proxied by Ethiopia_4500BP), southern African foragers (per South_Africa_1900BP), central rainforest Africans (per Mbuti), and eastern African pastoralists, represented by both ancient and present-day groups (Pastoral Neolithic [Kenya_PN] and Dinka), using Yoruba as a fixed first source representing West African-related ancestry (**Figure 1c and Supplementary Tables 4 and 5**). Among the 62 Iron Age and historical individuals from south central Africa, 13 cannot be statistically distinguished from forming a clade with Yoruba alone in qpWave (rank 0, p > 0.05). This pattern may reflect ancestry consistent with a single West African–related source relative to our curated reference set, or alternatively limited power due to relatively low SNP coverage and/or a low proportion of non–West African–related ancestry. Excluding individual I8379 from Kalala Island, where all one- and multi-source qpAdm models were rejected (p <0.05), we identified admixture models for 39 of the remaining ancient individuals. Most of them are consistent with two-way admixture, primarily involving West African-related ancestry combined with ancestry related to central African rainforest populations and to a lesser extent with southern African foragers. A subset of individuals required an additional eastern-related ancestry, including a Pastoral Neolithic-related component, consistent with a three-way admixture scenario (**Figure 1c, Supplementary Tables 4 and 5**). Nine individuals failed to be modeled even with a 3-way admixture model although they do not appear to be outliers in PCA (**Figure 1b**).

Using the software DATES^48^, we dated the admixture between Yoruba and central African rainforest populations (Mbuti) for a subset of individuals consistent with a two-way qpAdm model between these sources (**Supplementary Table 4**). These individuals were grouped according to radiocarbon age into Group1 (n = 4, range 1314-1633 calCE) and Group2 (n = 11, range 1639-1950 calCE). Group1 admixture time ranges from 118 calBCE to 1548 calCE and Group2 from 902 calBCE to 746 calCE. These estimates overlap the admixture time reported for present-day Zambians (range 410-942 CE)^22^. Taken together, the results are consistent with admixture between Bantu related populations and central African rainforest hunter gatherers having occurred well before the lifetimes of the individuals analyzed here, and they are most plausibly interpreted as reflecting event(s) that took place among the ancestors of Bantu related groups as they moved toward south central Africa.

The distinct position of Kalala Island individual I8379 in the PCA (**Figure 1b**) suggests an ancestry profile that differs from that of other individuals from south central Africa. This observation is consistent with previous findings in present-day BaTwa groups from Zambia^23^, including populations from the Kafue River region, which indicate the presence of a non-Bantu African ancestry component derived from a now-extinct forager population with no close affinity to present-day groups. To test formally whether ancestry related to LSA hunter-gatherers could account for this signal in I8379, we applied qpAdm using ancient south central African LSA individuals as a potential source population. Under this framework, the Kalala Island individual can be adequately modeled as a two-way admixture between Bantu-related ancestry (61.6%± 2.0%) and ancestry related to ancient LSA individuals from Fingira Rockshelter, Hora and Chencherere sites, Malawi^29,35^ (38.4% ± 1.9%). All alternative models substituting the south central African LSA source with ancient foragers from Kenya or Tanzania were rejected (p < 0.05, **Supplementary Table 6**). This interpretation is further corroborated by a qpAdm run in which the non-Bantu-related ancestry component is represented as a mixture of eastern, central African rainforest, and southern African-related ancestries previously identified among ancient south central African hunter-gatherers^29^. Under this framework, the I8379 individual can be adequately modeled as deriving 56.8% (± 4.3%) of their ancestry from West African-related sources, together with 16.0% (± 3.4%) eastern African hunter-gatherer-related, 17.1% (± 3.4%) central African rainforest hunter-gatherer-related, and 10.2% (± 1.6%) southern African hunter-gatherer-related ancestry. All intermediate models were rejected (p < 0.05; **Supplementary Table 7**).

We dated the admixture in this Kalala Island individual using Yoruba and a second source comprising ancient LSA foragers from Malawi, Kenya and Tanzania combined with present-day southern African foragers. This analysis yields a range of admixture time of 632-1438 calCE. For comparison, admixture dating of present-day BaTwa from the Kafue region performed in Breton et al.^23^ produced a more recent estimate, with an admixture time ranging from 1477 to 1566 CE. The difference in inferred admixture dates suggests continued or additional gene flow in the Kafue region after the period represented by the Kalala Island individual (1516-1667 calCE).

### Fine-scale population structure in south central Africa

As shown in the previous section, a large proportion of the genomes of ancient individuals from south central Africa (Iron Age and historical periods) is associated with Bantu-related ancestry, with minor but potentially meaningful pastoralist- and forager-related ancestry components. Given the clear evidence of mobility, contact, and population absorption in the linguistic and archaeological records^4,8–10,13,14,46,49,50^, it remains unclear whether fine-scale genetic substructure, potentially reflecting different migrations of distinct Bantu-speaking groups during or after the initial Bantu expansion, can be detected in this region.

To investigate such fine-scale population structure in south central Africa, we applied a suite of haplotype-based methods using genome-wide data from present-day and ancient individuals. (**Supplementary Table 8**). These analyses include clustering approaches using the Leiden algorithm^51^, FineSTRUCTURE^52^, and Total Variance Distance (TVD)^53,54^, combined with principal component analysis (PCA) of the ChromoPainter^53^ co-ancestry matrix. We first ran the Leiden algorithm on a dataset comprising 2,514 present-day African and non-African individuals, as well as 54 imputed ancient individuals from sub-Saharan Africa (**Supplementary Table 9**). From this analysis, we identified two clusters composed of individuals from south central Africa, including almost all Iron Age, historical, and present-day individuals from Malawi and Zambia, all individuals from the “Mozambique” group, and a large proportion of individuals from Tanzania (”Tanzania_TanzaniaMixed” group). Furthermore, the published ancient individuals Tanzania_Lindi (I14001.AG)^55^ and Tanzania_Pemba600BP (I2298.AG)^35^ also fall within these two clusters.

To further characterize this structure, we subsequently restricted our analysis to individuals found within these two clusters and applied FineSTRUCTURE (hereafter FS*_south central_*), PCA and population-based TVD, all of which corroborate the Leiden algorithm results (**Figure 2 and Supplementary Figures 1-5**). FS*_south central_*analysis inferred a larger number of clusters, up to K = 10, but cluster stability and number of individuals with a concordance rate > 0.9 decreased substantially for K larger than two (**Supplementary Figures 2-3**). To provide a statistical measure of genetic differentiation between population within clusters (with n>= 4) identified at K=2 from FS*_south central_* (**Supplementary Table 10** for the final clustering of modern south central Africans), we computed TVD based on the ChromoPainter painting profiles using either phased haplotype information (**Figure 2a,b and c**) or unphased genotype (“unlinked” mode of ChromoPainter, **Supplementary Figure 5**). The two clusters remain detectable using allele frequency patterns alone (unlinked mode), although the range of TVD values (0.0042-0.0052) is notably small. Moreover, despite the relatively low number of SNPs used in the analysis (∼170,000 SNPs), haplotype information further enhances resolution (global TVD range: 0.010- 0.22; cluster 1: 0.010-0.070; cluster 2: 0.021-0.15), revealing sharper fine-scale population structure and statistically significant differentiation (permutation test, p < 0.01, **Figure 2a**) within both clusters. Notably, Fwe and Shanjo groups from modern-day southwestern Zambia show the strongest genetic differentiation among cluster 2, likely reflecting recent isolation and/or strong founder effect as previously reported in Fortes-Lima et al.^22^ (**Figure 2c**). In addition, individuals from the Lozi ethnic group are distributed across both clusters, with 13 individuals assigned to cluster 1 and 10 to cluster 2 (**Figure 2d**).

**Figure 2.**
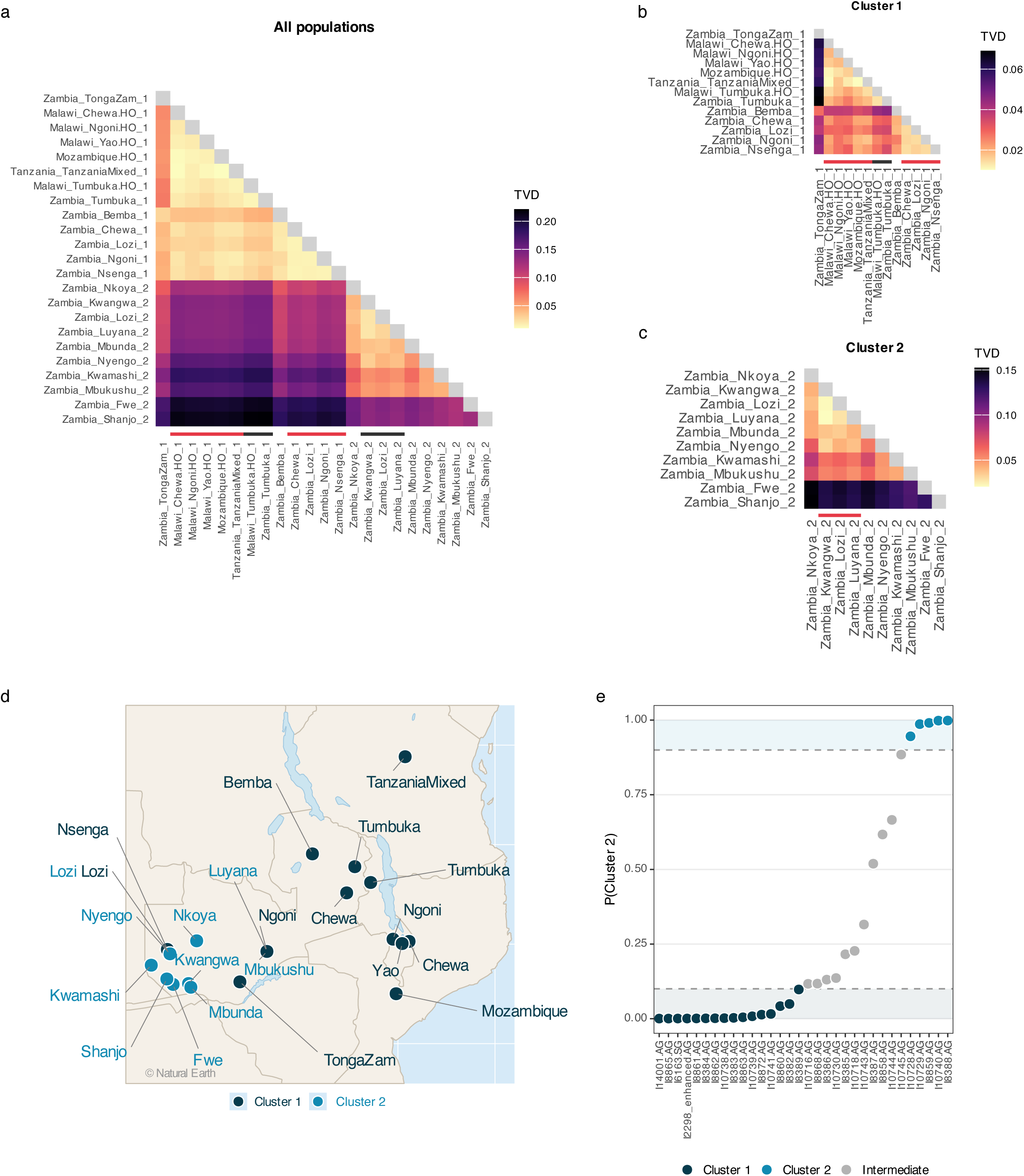
Haplotype-based clustering of modern Malawian and Zambian populations. (a) Pairwise Total Variation Distance (TVD) matrix across all populations, ordered by hierarchical clustering. Lower triangle shows TVD values (light yellow = low differentiation, dark purple = high differentiation); diagonal in grey. Horizontal bars below population labels on the x-axis indicate groups that could not be statistically distinguished by permutation test (p > 0.01); alternating red and black bars distinguish between independent non-differentiable groups. (b–c) TVD submatrices for Cluster 1 and Cluster 2 populations respectively, with independent color scales. (d) Geographic distribution of sampled populations colored by cluster assignment (Cluster 1: dark teal; Cluster 2: light blue). Some populations sharing the same sampling location appear as overlapping points. Note that Zambia Ngoni, Luyana, and Mbukushu were sampled in Lusaka and do not reflect their original geographic distribution. (e) Posterior probability of Cluster 2 membership for each ancient individual, estimated by Gaussian mixture model. Dashed lines indicate classification thresholds (p = 0.1 and p = 0.9). Points are colored by confident assignment: Cluster 1 (p(Cluster 2) ≤ 0.1), Cluster 2 (p(Cluster 2) ≥ 0.9), or Intermediate

Among the newly sequenced Iron Age and historical individuals, 14 were assigned to cluster 1 (as well as Tanzania_Pemba600BP and Tanzania_Lindi), five to cluster 2 with a posterior probability > 0.9 (**Figure 2e**), confirming a robust genetic division within south central Africa^21,22^. Three Kalala Island individuals with potentially overlapping chronologies mirror this pattern, with the I8382 individual (1667–1810 calCE) assigned to cluster 1 (posterior probability = 0.951), I8859 (1692–1950 calCE) to cluster 2 (posterior probability = 0.991), and I8858 (1893–1924 calCE) showing an intermediate position (posterior probability of cluster 2 = 0.617), suggesting that both genetic clusters coexisted at the same location, similar to what is observed among the modern Lozi. The temporal depth of this genetic structure is supported by ancient individuals from other sites: I10728 (Sinde Mission, where a different individual dates to 1639– 1797 CE) was confidently assigned to cluster 2 (posterior probability > 0.9), and I10745 (Ndonde, 1317–1400 calCE) also showed a strong affinity to cluster 2, albeit with a lower posterior probability (posterior probability = 0.885), indicating that this genetic structure was already established by at least the Late Iron Age, and potentially even earlier. Of note, the relatively low number of ancient individuals assigned to cluster 2 may partly reflect an archaeological bias toward the Lusaka-Livingstone areas of southern Zambia, and an absence of sites with human remains in westernmost Zambia (**Figure 1a**, **Supplementary Material Figure S1**). In addition, given the comparatively recent dates of individuals in cluster 2, it is possible that this cluster reflects multiple migration events, including some from the historical period (**Supplementary Table 1**).

### Different Bantu and non-Bantu-related affinities

Previous linguistic, archaeological, and historical studies^8,10,49,56^ have highlighted the complexity of the Bantu expansion, with evidence of layering, convergence, and back-migration across south central Africa. To evaluate whether the observed west-east genetic division within this region reflects differences in genetic proximity to Bantu and/or non-Bantu populations, we applied SOURCEFINDv2^57^ using present-day and ancient individuals from Clusters 1 and 2 as targets. SOURCEFIND identifies the source populations sharing the most recent common ancestry with each target; as a consequence, if the target population is genetically upstream of migration events, it is expected to copy predominantly from downstream groups. Importantly, SOURCEFIND inferences should be interpreted in relative rather than absolute terms: they reflect which source populations best match the haplotypic diversity of each target given the set of available donors and sources, but do not imply direct descent from those sources. We therefore performed two separate runs of ChromoPainter and SOURCEFIND, each using a different set of donor and source populations (**Supplementary Tables 11 and 12**). The first run included present-day Bantu-related groups from across sub-Saharan Africa, in addition to non-African and non-Bantu populations, while the second was restricted to Bantu-related populations thought to be upstream of south central Africa^8,21,22,43^, namely from Nigeria, Cameroon, Angola, the Republic of Congo and the Democratic Republic of Congo (DRC) in addition to non-African and non-Bantu populations.

In the first run, the predominant best sources for groups within cluster 1, including ancient individuals (IA_Historical_1), were the south central DRC Luba-Lulua group, the Mongo/Nkundo equatorial DRC group and mainly neighboring southeastern Bantu-related populations such as Mozambique Tsonga, South Africa Venda and Tsonga as well as Zimbabwe. In contrast, groups within cluster 2 were best matched to Bantu-related populations from the Bié plateau of Angola (Ovimbundu) and from Namibia (Wambo/Ovambo) (**Figure 3a, Supplementary Table 11**). The Zambian Lozi ethnic group, among whom 13 of 23 individuals were assigned to cluster 1 and 10 to cluster 2, further confirmed these results: Lozi individuals in cluster 1 showed predominantly south central DRC-related ancestry, whereas those in cluster 2 exhibited higher levels of Angolan-related ancestry. To formally quantify this separation, we trained a random forest classifier using per-individual SOURCEFIND ancestry proportions from the first iteration (including all donors and sources) as predictors. The model achieved near-perfect classification accuracy (98.8%), with perfect sensitivity for cluster 1 (specificity=1.00, sensitivity=1.00) and high sensitivity for cluster 2 (specificity=0.977, sensitivity=1.00). Only three of 251 individuals were misclassified, all from cluster 2, and none from cluster 1. The most informative predictors were Angola_Ovimbundu and groups from the Great Lakes region (DRC_Shi, Uganda_Kiga and Uganda_Nkore) (**Supplementary Figures 9 and 10**). Predicted probabilities per population were highly consistent within each cluster, with nearly all populations clustering tightly near 0% or 100% probability of cluster 2 assignment, confirming that the two clusters represent a coherent ancestry distinction across Iron Age, historical and modern individuals (**Supplementary Figure 11**).

**Figure 3.**
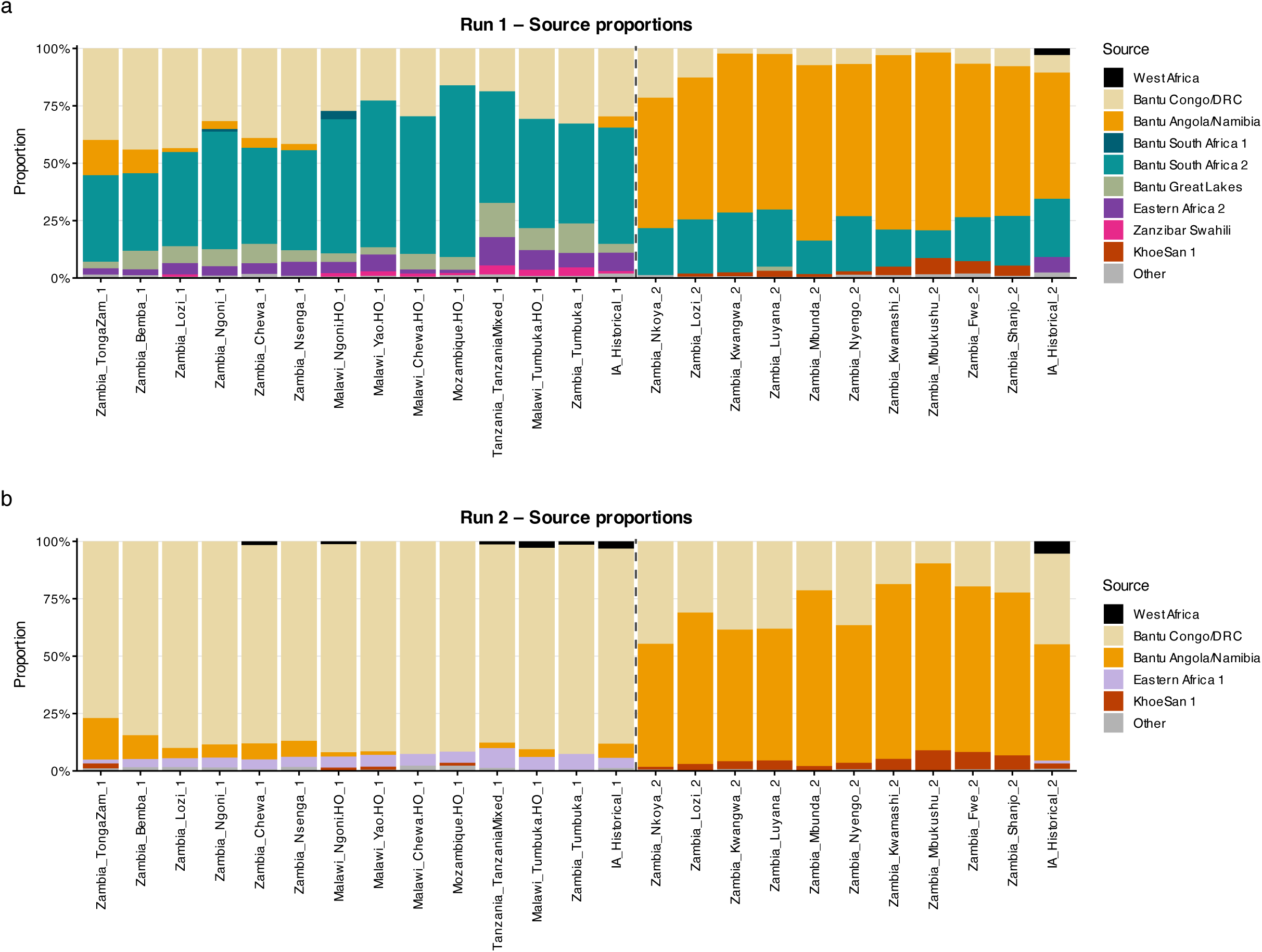
SOURCEFIND ancestry proportions in modern and ancient Malawian and Zambian populations. Stacked bar charts showing the average proportion of ancestry derived from each source population group for Run 1 **(a)** and Run 2 **(b)**. Source populations were aggregated into metaclusters based on the first two levels of the Leiden clustering (**Supplementary Table 9).** Source populations contributing less than 1% of ancestry on average were collapsed into the “Other” category. IA_Historical_1 and IA_Historical_2 denote Iron Age and historical ancient individuals assigned to Cluster 1 and Cluster 2, respectively with posterior probability > 0.9. 2-level cluster labels correspond to the following source population groups: 1_1 = Bantu Angola/Namibia, 1_2 = Bantu Congo/DRC, 1_3 = West Africa, 2_1 = Eastern Africa 1, 2_2 = Eastern Africa 2, 2_3 = Bantu Great Lakes, 3_3 = Zanzibar Swahili, 4_1 = Bantu South Africa 1, 4_2 = Bantu South Africa 2, 5_1 = KhoeSan 1, 5_4 = KhoeSan 2.

In the second run, restricted to Bantu-related populations upstream of south central Africa, the Angola/Namibia signal for cluster 2 was preserved, with Ovimbundu and Wambo remaining the dominant sources across all cluster 2 populations (**Figure 3b, Supplementary Table 12**). For cluster 1, DRC Luba-Lulua emerged as the predominant source in the absence of southeastern Bantu donors. Interestingly, cluster 2 populations also showed an increased contribution of DRC Luba-Lulua compared to the first run, suggesting that the South African Bantu-related ancestry observed in the first run is at least partly replaced by Luba-Lulua ancestry when southeastern donors are unavailable. This pattern is consistent with a scenario in which the southern African Bantu-related ancestry reflects a downstream derivative of a south central DRC-related ancestry component.

FastGLOBETROTTER^58^ analyses, restricted to the same upstream Bantu-related donor populations as the second SOURCEFIND run, detected admixture signals in several target groups from both clusters (**Supplementary Table 13**). In all cases, the major source (S2, ∼94–97%) was highly consistent with SOURCEFIND results: cluster 1 groups were best modelled as deriving predominantly from a DRC Luba-Lulua-related source (S2). Cluster 2 groups were best modelled with Angola Ovimbundu as the dominant source, with Namibia Wambo and DRC Luba-Lulua as additional contributors, again consistent with the SOURCEFIND results. The minor source (S1, ∼4–6%) revealed a distinct signal in each cluster: cluster 1 groups showed affinities to eastern African populations including Somali, Sandawe and Ethiopian groups, while cluster 2 groups were best matched to KhoeSan-related populations such as Xuun, Khwe and Ju hoan. Although these non-Bantu ancestry components are consistently detected across groups, their low proportions (∼5%) could reflect (i) genuine low-level admixture with non-Bantu populations and/or (ii) ancestry mediated through additional Bantu migrations that simultaneously carried and introduced non-Bantu-related ancestry. Given the high level of genetic similarity between Bantu groups, the absence of detected admixture between distinct Bantu sources may reflect limited statistical power rather than a true absence of such events.

Previous studies^21,43^ proposed a serial founder model for the Bantu dispersal into southeastern Africa, with successive movements from northern Angola into Zambia, then Malawi and northern Mozambique, and further south. The strong affinity of cluster 2 to Angolan-related populations, suggests that cluster 2 shares more recent common ancestry with Angolan/Namibian Bantu-related groups than cluster 1 does, possibly reflecting a closer relationship to a shared ancestral population in the western/southwestern part of the Congo basin or Angola region (although Ovimbundu and Wambo should be considered the closest available proxies rather than direct sources). However, the serial founder model would predict that cluster 1, as a downstream derivative of the same dispersal, should similarly show Angolan-related ancestry when southeastern donors are excluded. This is not what we observe: cluster 1 is instead overwhelmingly best matched to present-day south central DRC (Luba-Lulua)-related ancestry, suggesting a distinct and independent migration.

To further evaluate this, we performed identity-by-descent (IBD) analyses, which measure recent relatedness from shared genomic segments (≥4 cM). Using Ovimbundu as the reference Angolan population, the serial founder model makes three specific predictions: (i) cluster 2 should show higher IBD with Ovimbundu than cluster 1 does, reflecting its closer proximity to the proposed Angolan source; (ii) cluster 1 and cluster 2 should share more IBD with each other than cluster 2 shares with Ovimbundu, given their proposed ancestor-descendant relationship; and (iii) cluster 1 should show stronger IBD with Ovimbundu than with south central DRC populations (Luba Lulua), reflecting passage through the Angolan corridor. The first prediction is confirmed, while predictions (ii) and (iii) are rejected. Cluster 2 shares on average substantially more IBD with Ovimbundu than cluster 1 does (2.62 vs 0.64 cM, diff = +1.98 cM [95% CI: 1.55–2.38], one-sided p.val = 0.0015), confirming that the Angolan affinity is specific to cluster 2. However, cluster 2 shares more IBD with Ovimbundu than with cluster 1 (2.62 vs 1.65 cM, diff = +0.96 cM [95% CI: 0.59–1.36], one-sided p.val = 0.0015), indicating that cluster 2 and Ovimbundu groups are more recently related than cluster 2 and cluster 1 are. Finally, contrary to prediction (iii), cluster 1 shows significantly more IBD with DRC Luba-Lulua than with Ovimbundu groups (0.95 vs 0.64 cM, diff = +0.30 cM [95% CI: 0.030–0.566], one-sided p.val = 0.020), inconsistent with cluster 1 having passed through the same Angolan corridor.

Taken together, our SOURCEFIND, FastGLOBETROTTER and IBD analyses consistently reject a simple serial founder model and point instead to at least two distinct Bantu-related dispersals, correlating with layered patterns of linguistic differentiation in southern DRC and convergence in Zambia, as discussed below. Our results are more consistent with the model proposed by Fortes-Lima et al.^22^, in which the DRC represents a major crossroads for the expansion of Bantu-speaking populations, giving rise to multiple genetically distinguishable dispersal routes rather than a single linear movement. We therefore propose that the two clusters reflect, at least, two such distinct dispersals that reached present-day Zambia and Malawi through different routes and/or at different times. One possible interpretation is that cluster 1 and cluster 2 share a common ancestral population in the south central DRC region, but that the ancestors of cluster 2 subsequently moved through or had prolonged contact with southwestern DRC or Angolan populations before or after reaching Zambia, explaining both the shared DRC-related background and the strong Angolan affinity specific to cluster 2.

### DRC and south central Africa in the context of the Bantu expansion

To further characterize the relationships between the two south central clusters and other Bantu-related populations, we performed an additional ChromoPainter/SOURCEFIND analysis in which cluster 1 and cluster 2 were included as sources alongside the other groups used in the previous section (run 2), allowing us to assess their relative contributions to the ancestry of Bantu-related target groups from Angola/Namibia, southern, and Great Lakes region of Africa (**Figure 4a, Supplementary Table 14**). Cluster 1 emerges as the almost unique Bantu-related source for South Bantu target populations, contributing 76–100% of ancestry across all groups. For Great Lakes groups, cluster 1 again contributes substantially (31–61%) but not exclusively. Cluster 2 shows no signal in southern Bantu or Great Lakes targets but is present in Angola/Namibia, where it appears as the dominant Bantu source (29–48%); however, several DRC groups also contribute substantially to this region (11–59%), including DRC_MongoNkundo (up to 69%), DRC_Pende (up to 27%), DRC_Ndibu (up to 13%), and DRC_Mbala (up to 14%).

**Figure 4.**
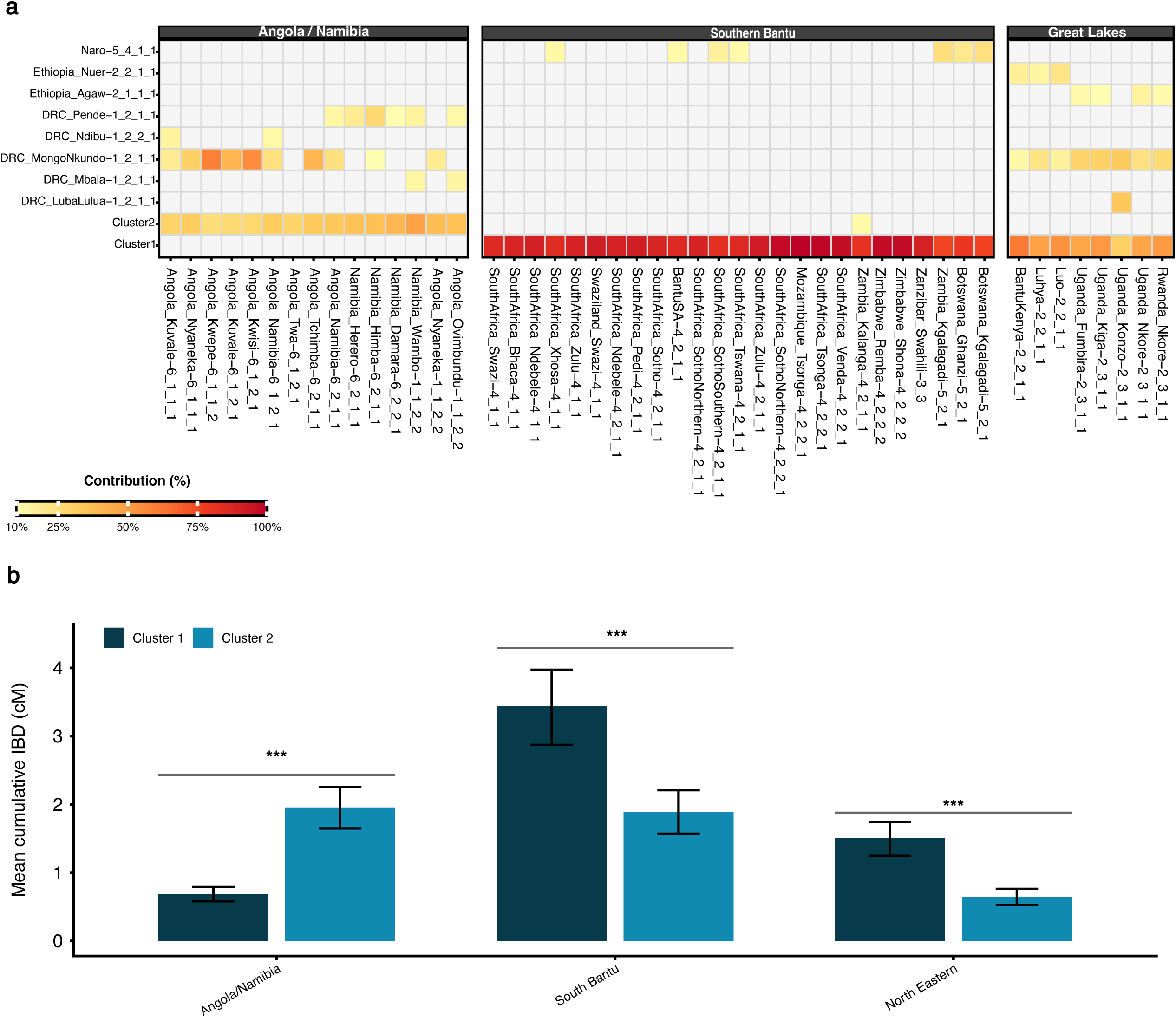
South-central African ancestry (Cluster 1 and Cluster 2) in modern Bantu-speaking populations across sub-Saharan Africa. **(a)** Heatmap showing sources contributing more than 10% to each target population (inferred with SOURCEFIND), grouped into Angola/Namibia, South Bantu, and Northeastern geographic regions. Color intensity reflects the percentage contribution of each source population (light yellow = 10%, dark red = 100%). Only sources exceeding 10% in at least one target population are shown. **(b)** Mean cumulative identity-by-descent (IBD) sharing (≥4 cM segments) between Cluster 1 (dark blue), Cluster 2 (light blue) and three geographic regions of modern Bantu-speaking populations. IBD values are averaged across all possible individual pairs within each population group, including pairs with zero IBD sharing. Error bars represent 95% bootstrap confidence intervals (n=1,000 genomic block resampling). Significance brackets indicate BH-corrected p-values from two-sided tests (*** p < 0.001)

We complemented these SOURCEFIND results with IBD analyses. The IBD results corroborate the geographic specificity of the two clusters: cluster 2 shares significantly more IBD with Angola/Namibia populations than cluster 1 (average IBD = 1.95 vs 0.69 cM, two-sided p < 0.001), while cluster 1 shares significantly more IBD with both southern Bantu (3.44 vs 1.89 cM, p < 0.001) and Great Lakes populations (1.50 vs 0.64 cM, p < 0.001; **Figure 4b, Supplementary Figure 12**). Importantly, the IBD signal of cluster 2 with Angola/Namibia (1.95 cM, 95%CI [1.65-2.25]) stands clearly above that of all central DRC populations (min=0.41cM, max=0.76 cM; **Figure 5a**), demonstrating that its affinity with Angola/Namibia is specific and not simply a reflection of general Bantu-related background relatedness. Similarly, IBD sharing between cluster 1 and southern Bantu populations (3.44 cM, 95%CI [2.87-3.97]) exceeds that of all central DRC populations, with DRC_LubaLulua being the closest DRC group (0.95 cM, 95%CI [0.75-1.18]) but still far below cluster 1. Finally, examining IBD between central DRC populations and each south central African cluster (**Figure 5b**), cluster 2 shows systematically higher IBD sharing than cluster 1 across many DRC populations, with significant differences observed for DRC_Ding, DRC_Mbala, DRC_Mbuun, DRC_Ngwi, and DRC_Pende (p < 0.002). This pattern may reflect more recent connections between cluster 2 and DRC groups relative to cluster 1, or alternatively, that the DRC population(s) at the origin of cluster 2 underwent a stronger founder effect, inflating IBD sharing through drift compared to the source population(s) of cluster 1.

**Figure 5.**
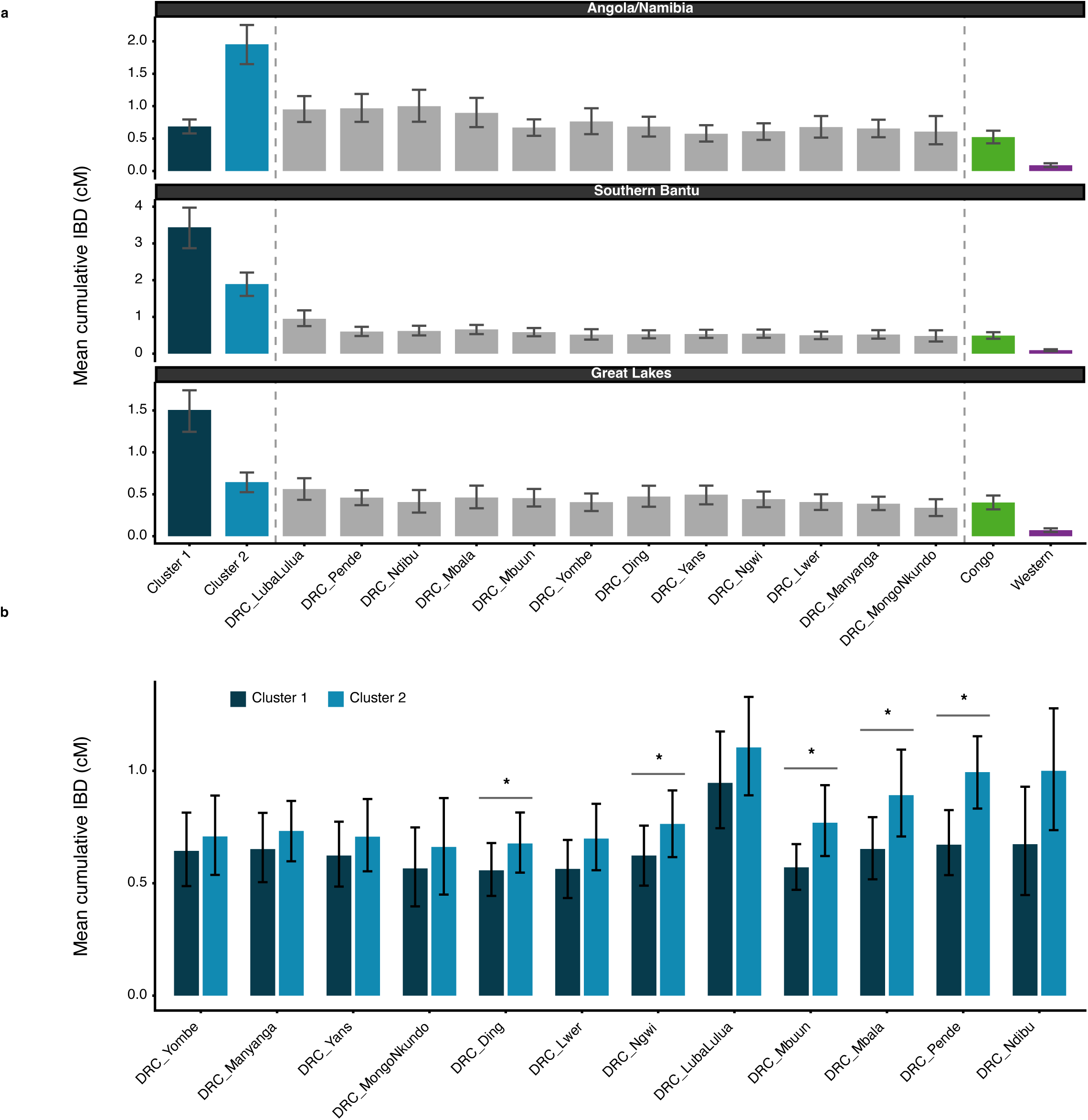
IBD sharing between modern Bantu-related populations from different regions of sub-Saharan Africa, south central African clusters, and DRCpopulations. **(a)** Mean cumulative IBD sharing between Angola/Namibia, southern and Great Lakes regions and south central cluster 1, cluster 2 (only present-day individuals), populations from DRC, pooled Congo populations, and pooled Western African populations (Yoruba and Ibibio). Error bars represent 95% bootstrap confidence intervals (n=1,000 genomic block resampling). **(b)** Mean cumulative IBD sharing between each DRC population and Cluster 1 (dark blue) versus Cluster 2 (light blue). Error bars represent 95% bootstrap confidence intervals (n=1,000). Significance brackets are shown only for populations with statistically significant differences between clusters (two-sided bootstrap test, BH-corrected; *: p < 0.003).

Altogether, these results provide insight into the dispersal histories connecting south central Africa to broader Bantu-related populations. For southern Bantu, the pattern is consistent with a southward expansion from south central DRC populations (Kasaï region), representing an ancestral lineage that gave cluster 1, which then expanded further south. This is supported both by the near-exclusive contribution of cluster 1 to southern Bantu ancestry (76–100%) and by its substantially elevated IBD sharing with these populations (average = 3.44 cM) relative to any DRC group (max 1.17 cM for DRC_LubaLulua). For Great Lakes populations, the picture is more complex. Although cluster 1 contributes substantially (31–61%), these populations also show meaningful ancestry from other DRC groups. We note that the specific identity of these DRC sources should be interpreted with caution. However, unlike southern Bantu-related groups, which derive almost exclusively from cluster 1, Great Lakes populations retain ancestry from additional DRC-related lineages, suggesting that their ancestry is not fully captured by cluster 1 alone. This points to a scenario in which cluster 1 and Great Lakes Bantu groups share a common ancestral population, plausibly located in south central or southeast DRC, from which they diverged: one lineage expanding southward to give rise to cluster 1 and ultimately southern Bantu, and another moving toward the Great Lakes region. Alternatively, this region may have been settled first by a different Bantu-related lineage, and present-day Great Lakes Bantu groups may result from later mixing with migrants related to cluster 1. Cluster 2, by contrast, shows no signal in either southern Bantu or Great Lakes targets but emerges as the dominant Bantu source for Angola/Namibia groups, pointing to a distinct and geographically separate dispersal trajectory consistent with the results of the previous section.

## Discussion

This study demonstrates the value of legacy archaeological collections, which for many sites in this study were not previously analyzed or dated (**Supplementary Material**). Even for the best-documented sites in our dataset, newly generated direct dates will force a reassessment of chronologies long based on material culture and/or indirect, pre-AMS dates on charcoal. With rare exceptions, the sampled human remains have not been subject to osteological analysis, and the archival and genetic research brought together here points to future lines of bioarchaeological inquiry on individual life histories and population-level dynamics during the Iron Age. This contributes to a recent surge of interest in the Iron Age archaeology of south central Africa, after decades of dormancy^4,5,50,59–61^.

Our genetic results suggest that the dominant Bantu-related ancestry component in populations from the eastern part of south central Africa is most closely represented, in our reference panel, by Luba-Lulua groups from the south central DRC. The historically documented Luba-Lunda states are relatively recent political formations, traditionally dated to around the 16th century CE onward, yet our analyses identify a Luba-related group as the best genetic proxies for the dominant Bantu-related ancestry even in earlier individuals in our study (cluster 1). Archaeological and linguistic evidence places the first Bantu-speaking communities in parts of Zambia and Malawi by the mid first millennium CE ^4,9,49,50,62^, centuries before the Luba-Lunda states formed, creating a temporal gap that raises the possibility of multiple migrations stemming from a shared region within DRC.

In this scenario of a “spread-over-spread” model, the initial Bantu-speaking populations reaching Malawi and Zambia may have already descended from a lineage closely related to present-day Luba groups. Under this view, the Luba state represents a relatively recent political crystallization within a much older and geographically widespread demographic continuum. The Upemba Depression of southeastern DRC provides a plausible archaeological context for this continuum. Indeed, archaeological studies report evidence of trade connections between the Upemba Depression and the Zambian Copperbelt region centuries before the emergence of the Luba state, and further suggest that the Luba people themselves originated in this region^15,16^, making the Upemba Depression a plausible demographic hub ancestral to cluster 1 and a promising area for future ancient DNA research. Although our data cannot definitively confirm this scenario, the widespread Luba-related signal in cluster 1 and the distinct Angolan-related ancestry characterizing cluster 2 point toward at least two independent waves of Bantu-speaking populations radiating from a DRC core, with cluster 1 likely representing part of an expansion route continuing southward toward present-day southern Africa, likely as an equally complex layered expansion history^49^. This supports neither a simple wave of advance nor a serial founder model, but rather a more complex layered history of movements where south central Africa acts both as a crossroads and a point of divergence between expansion routes.

The northeastern half of Angola, western Zambia, and southern DRC (from Kananga east to Lake Tanganyika and south to the border with Zambia) represent important corridors of expansion and migration in the linguistic and historical records^8,10,17,18^, yet remain poorly represented archaeologically and by modern and ancient genetic data. After decades of research disentangling later layering from shared ancestral origins, a broad consensus is emerging^10^ pointing to the savannas south of the equatorial forests as a region of particularly complex language spread, convergence, and likely widespread multilingualism^9,10,14,49,56^. This research indicates that the expansion of Bantu languages into south central Africa unfolded in successive waves. The first involved the Kongo Cluster spreading across the Kwango River into west central Africa, followed by the Lweta or southwest Bantu group moving into western Zambia, Angola, and northern Namibia, and then the Central Savanna Group (including Luban languages) spreading across southern DRC and into northern Zambia. A subsequent stream entered Zambia from the southern shores of Lake Tanganyika, represented by the Sabi and Botatwe language groups, which eventually spread across eastern, central, southern, and western Zambia and beyond the Zambezi. Further splits from the main Bantu trunk carried languages northward into the Great Lakes region, where they may have encountered communities descending from earlier rainforest dispersals. A final major divergence produced the communities responsible for spreading Bantu languages along the eastern corridor from the Swahili coast through Tanzania, Malawi, and Mozambique, and ultimately into Zimbabwe, Botswana, and South Africa. Languages of this last eastern group may have spread across eastern and southern Zambia before being overlaid by languages of the Sabi and Botatwe groups^9,49^ but also spread into these parts of Zambia from Malawi and Zimbabwe in more recent centuries. Precolonial polities, including the Lozi, Mutapa, and major west-central African states, and later migrations such as the 19th- century Mfecane and subsequent European colonization, created dynamic landscapes in which long-distance mobility was common. Groups such as the Ngoni, Yao, and Chokwe, all found in Malawi and Zambia today, expanded rapidly into the region in the 19^th^ century, incorporating diverse local communities and potentially reshaping regional genetic landscapes in the process^4,17,18,63–66^.

When genetic, archaeological, linguistic, and historical evidence are considered together, it would be overly simplistic to treat early arriving Bantu communities as static. The genetic structure observed today likely reflects repeated episodes of population movement, political centralization, and long-distance trade over the past one to two millennia. We identify four episodes in which the genetic and linguistic evidence attest to parallel processes of layering and mixing, to better appreciate the value of haplotype-based analysis generating fine-scale population structure in correlating with linguistic and archaeological data.

On the narrowest geographical and chronological scale, the unique features of the Lozi, Shanjo, and Fwe populations vis-à-vis clusters 1 and 2 correlate with linguistic data. The mix of cluster 1 and 2 individuals in the Lozi population mirrors the linguistic mix of Luyana (more closely related linguistically with southwestern African groups associated with cluster 2 and associated with the earliest Barotse floodplain polity) and southeast Bantu (more closely related linguistically with Narrow East Bantu language speaking groups, here correlated with cluster 1)^67^. This population and language mixing resulted from the Mfecane, in which displaced populations migrated north from what is now northeast South Africa, crossing the Zambezi River and seeking to overtake a Barotse floodplain polity. That some individuals maintained one or another genetic identity likely reflects the fractured politics of the indigenous Luyana and foreign Makolo competing for control of the floodplain polity and of central and western Zambia more generally^65^. Similarly, the strong outlier signal connecting Shanjo and Fwe populations, in contrast to others in cluster 2, correlates to their position as an independent subgroup of western Botatwe with a unique linguistic ancestor that marks them as more closely related to each other than to any other Bantu languages, perhaps linked to fleeing Luyana expansion before the 19^th^ century^56^. Like genetic ties to Ovumbundu and Ovambo populations in western Zambia, influence from Southwest Bantu languages carried into western Zambia through trade (including in slaves) from the 16^th^ or 17^th^ century, perhaps accounting for the mixed affiliations of Kalala Island individuals^18,68^.

This same geography of influence can be seen earlier, through the late first millennium spread of the so-called matrilineal belt from southwest Angola communities across Zambia to Malawi and beyond ^9,69^. This west-to-east introduction of kinship practices, which emphasize matrilineal ties in controlling marriage and procreation, would leave a genetic signature. Linguistic evidence suggests the widespread adoption of these practices from Angolan groups into the middle Kafue at the turn of the first millennium, with a rapid expansion of speakers of Kafue (Botatwe) languages across the Batoka and to the east and northeast of the middle Kafue through matrilineal ties starting from the thirteenth century.The Ndonde (I10745) individual’s close ties to cluster 2 might suggest the introduction of this marriage pattern involved both borrowing the practice and incorporating outsiders who introduced it.

At the broadest scale, the relative relatedness of cluster 1 individuals to southeastern, Great Lakes, and Luba-Lulua populations corresponds well to the linguistic evidence for the convergence in southern DRC and Zambia of populations speaking at least four major branches of Bantu that developed after Southwestern Bantu languages (associated with cluster 2) split off: Central Savanna (Luban), Maniema, Central Woodlands (Sabi/Botatwe), and West Tanzania/Southeast Bantu (Great Lakes split off between Central Woodlands and West Tanzania/Southeast Bantu)^8,10,49^. For the historical reasons outlined earlier, speakers of descendant language of these four groups continued to have contact and converge in southern DRC and across south central Africa for over a millennium.

In conlusion and consistent with previous work^20,21^, individuals from Zambia and Malawi across the Iron Age, historical, and present-day periods carry relatively modest proportions of non-Bantu ancestry. The ancient outlier individual from Kalala Island (I8379), who exhibits a substantial proportion of LSA forager-related ancestry, demonstrates that pockets of local forager-related ancestry persisted well after Bantu-related groups became regionally dominant. Taken together, these findings indicate: (i) the identification of two distinct Bantu-related ancestry components, (ii) the south central DRC-related signal dominating in part of Malawi and Zambia, (iii) the Angolan-related component in western Zambia, and (iv) the generally low but variable levels of non-Bantu ancestry are broadly consistent across the Iron Age, historical, and present-day individuals in our study, though their resolution remains tied to currently available reference panels. Incorporating additional populations, particularly from under sampled regions of the DRC and eastern Angola (e.g., Katanga, Upemba regions), will likely reveal finer-grained substructure within the broad south central DRC and Angolan-related components identified here. Nevertheless, our results suggest that haplotype-based methods can be usefully applied to Iron Age African genomes to investigate the Bantu expansion, even with current reference panel limitations. Future studies combining higher-coverage ancient genomes with broader modern reference panels will help further refine our understanding of these complex demographic histories.

## Methods

### Skeletal sampling and radiocarbon dating

For sites curated at the Livingstone Museum (LM), Zambia (all Zambian sites plus most of Nkudzi Bay), human remains were inventoried, and archival and bioarchaeological analysis enabled the differentiation of individuals in fragmentary and commingled collections, to avoid double-sampling. Skeletal samples maximized chances of DNA success and minimized impacts on collections, prioritizing petrous and tooth, then postcranial bone, and seeking isolated, fragmented, or otherwise less informative remains. Digital and photographic records of sampling activity were deposited with the museum. All LM samples that produced readable aDNA, plus two MNR samples, were submitted to the Pennsylvania State University Radiocarbon Laboratory for radiocarbon dating via accelerator mass spectrometry (AMS). Collagen carbon and nitrogen concentrations and stable isotope ratios were measured at the Yale Analytical and Stable Isotope Center. We calibrated all dates using OxCal v.4.4^70^ with the SHCal20 curve^71^.

### Library preparation

For each individual, we prepared double- and/or single-stranded libraries treated with partial uracil-DNA glycosylase (UDG) to reduce characteristic ancient DNA damage (**Supplementary Table 1**). We then performed in-solution enrichment targeting >1.2 million genome-wide SNPs^45,72^ and sequenced the enriched libraries on Illumina platforms. We processed raw data using standard ancient DNA bioinformatic pipelines and applied additional SNP filters based on SNPs being part of a compatibility SNP set^73^ designed to minimize biases in downstream population genetic analyses combining modern and ancient individuals.

### Datasets

Ancient 1240K pseudo-haploid data for Iron Age and historical individuals from south central Africa were merged with a reference panel comprising published ancient 1240K and present-day whole-genome sequence data from African and non-African populations. SNPs identified as problematic when co-analyzing 1240K capture and shotgun sequencing data^73^ were excluded, resulting in a dataset of 656,883 SNPs. Ancient individuals with less than 15,000 SNPs covered on 1240K or less than 10,000 SNPs covered on the compatibility snpset were excluded (**Supplementary Table 1**)

To investigate fine-scale population structure in south central Africa, we constructed a separate dataset by merging Human Origins genotyping array data (HO panel) from African and non-African individuals with African populations from the African Neo panel^22^. We retained bi-allelic SNPs present in both panels and merged the datasets using PLINK v1.9 (https://www.cog-genomics.org/plink/). We then applied the following quality control filters sequentially: (i) removal of individuals with more than 95% missing data; (ii) removal of cryptically related individuals up to third-degree relatedness using KING v2.3.2^74^; (iii) removal of bi-allelic SNPs with more than 10% missingness; (iv) removal of SNPs deviating from Hardy-Weinberg equilibrium within each population (n ≥ 10) using PLINK v1.9; and (v) removal of individuals with excess or deficit heterozygosity, defined as ± 3 standard deviations from the population mean. The dataset comprised 167,912 segregating bi-allelic SNPs.

### Imputation of ancient individuals and phasing

The 54 ancient sub-Saharan Africans, with more than 1X coverage on 1240K SNP set were imputed using GLIMPSEv1.0.0^75^ and 1K genome phase 3 as reference panel following the procedure described in Akbari et al.^76^. Following imputation, genotype probabilities were filtered per sample, retaining only genotypes with a posterior probability ≥ 0.80. The imputed ancient dataset was then intersected with the present-day reference panel (African Neo panel merged with HO panel) to retain only variants common to both datasets. The merged dataset was subsequently phased using SHAPEIT5^77^ using 1K genome phase 3 as reference panel.

For haplotype-based analyses, a subset of this phased dataset was used due to computational constraints. We retained populations with at least five individuals, with two exceptions: (i) all present-day individuals from Malawi and Zambia were retained regardless of population size, and (ii) for non-African, non-Bantu, and West African groups, only a representative subset of populations was included. The list of retained populations is provided in **Supplementary Table 8**

### PCA, qpWave, qpAdm

PCA were conducted using smartpca^78^ with the lsqproject and newshrink options using the dataset 1240K intersected with shotgun data (656,883 SNPs). All ancient individuals were projected on the PC axis computed using only modern individuals.

qpWave and qpAdm were conducted using ADMIXTOOLS^46^ with the following arguments: ‘inbreed: NO’ and ‘allsnps: NO’. We used the following set of right populations: Lemande, Mbuti, Dinka, Biaka, Bulala, Khomani_San, Russia_UstIshim, Israel_C and French. The source populations or left set of populations: Yoruba, Mbuti, Dinka, South_Africa_1900BP, Kenya_PastoralN, Ethiopia_4500BP. When “Mbuti” and “Dinka” were tested as left population they were removed from the right list.

For the ancestry deconvolution of ancient individual I8379, we used as right populations: Chimp, Lemande, Biaka, Khomani_San and French. The set of left population was composed of: Yoruba, Mbuti, South_Africa_1900BP, and Ethiopia_4500BP.

### DATES

DATES^48^ was run using maxdis=0.2 and binsize=0.001. To estimate the timing of admixture in calendar years, the DATES estimate (in generations) was converted to years using a generation time of 28 years. For the two groups, the ^14^C date range was defined as the minimum lower bound and maximum upper bound of the calibrated ¹□C confidence intervals across all individuals within groups. The admixture date range was then obtained by combining this ¹□C range with the 95% confidence interval of the DATES estimate (mean ± 1.96 SE): the earliest admixture bound was calculated using the oldest possible ¹□C date and the maximum number of generations, while the latest admixture bound used the most recent ¹□C date and the minimum number of generations.

### ChromoPainter

Unless otherwise stated, each individual was set to both donor and recipient, meaning that all individuals can copy from all others excluding themselves (-a mode). We first estimated global mutation probability and switch rate using 10 expectation-maximization iterations (-in -iM arguments) by averaging estimates obtained for chromosomes 1, 8, 15, and 22 (weighted by number of SNPs per chromosome). We then ran Chromopainter v2.0^52^ again using the same mode but fixing the two parameters to the estimated values. To complement haplotype-based analyses, we additionally ran Chromopainter in unlinked mode (-u mode), in which each SNP is painted independently without using haplotype phase information, providing a coancestry matrix that captures allele sharing patterns used for TVD comparisons.

### Leiden clustering

To identify broad genetic clusters, we applied the Leiden community detection algorithm to the Chromopainter coancestry matrix (chunckcounts.out) using the igraph R package^79^. The matrix was first symmetrized and sparsified by retaining for each individual only their top k connections (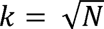, where N is the total number of individuals). To ensure graph symmetry, if individual A retained individual B among its top k connections but B did not retain A, the edge was preserved in both directions by taking the element-wise maximum between the matrix and its transpose. The resulting weighted network was then converted to a graph and partitioned using a recursive implementation of the Leiden algorithm with modularity as the objective function (resolution = 0.5, minimum community size = 20, maximum recursion depth = 4). At each recursion level, the algorithm was run 100 times and the partition maximizing modularity was retained. Each identified community was then treated as an independent subgraph and re-partitioned in subsequent iterations, allowing the detection of finer-grained structure nested within broader clusters. Recursion stopped when either the maximum depth was reached, a community could not be further subdivided, or a community fell below the minimum size threshold. Each individual was assigned a population-level cluster label combining their population name and cluster identifier on the form <*population_label*>_<*cluster_ID*>, and population-cluster combinations represented by fewer than four individuals were excluded from downstream analyses (**Supplementary Table 9**).

### Haplotype-based PCA

To compare pattern of haplotype-sharing inferred by CHROMOPAINTER we computed principal component analysis using the script provided in https://people.maths.bris.ac.uk/~madjl/finestructure-old/finestructureR.html and the ChromoPainter chunckcounts.out coancestry matrix.

### fineSTRUCTURE clustering

We clustered present-day south central African individuals using fineSTRUCTURE^52^ (FS_south central_). Specifically, we included present-day individuals assigned to Leiden clusters ‘1_1_1_1’ and ‘1_1_2_1’ in the Leiden algorithm described above. For this analysis, Chromopainter was run in all-versus-all mode (-a flag) using only these present-day individuals, and the resulting coancestry matrix (chunckcounts.out) was used as input for fineSTRUCTURE. We ran fineSTRUCTURE as in [refs ^41,53^]. We computed 2,000,000 iterations of the Markov Chain Monte Carlo (MCMC) algorithm, recording inferred clustering every 10,000 iterations. We then identified the single sampled clustering with the highest overall posterior probability. Starting from this clustering, we conducted an additional 100,000 hill-climbing steps to reach a nearby state with an even higher posterior probability. We ran the algorithm ten times starting from different seeds and selected the run with the highest posterior probability. We evaluated the accuracy of the clustering based on concordance rate per individual, similar to Leslie et al.^53^. As with the Leiden algorithm, each individual was assigned a population-level cluster label combining their population name and cluster identifier (<*population_label*>_<*FS_cluster_ID_*>), and population-cluster combinations represented by fewer than four individuals were excluded from downstream analyses.

### TVD

To quantify genetic differentiation between population-level clusters identified by FS*_south central_*, we computed pairwise Total Variation Distance (TVD) based on the Chromopainter chunk length coancestry matrix. For each individual, the coancestry vector was row-normalized to sum to one, representing the proportion of the genome copied from each donor individual. Individual copying vectors were then averaged within each population-cluster group to obtain a mean copying profile, and pairwise TVD between groups A and B was

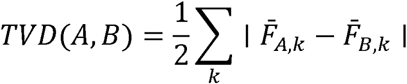

Where 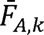, and 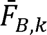 are the mean proportions of genome copied from donor group k by individuals in groups A and B respectively.

To assess the statistical significance of pairwise TVD values, we built a hierarchical tree by iteratively merging the pair of groups with the smallest TVD at each step and recomputing mean copying profiles after each merge, inspired by the procedure described in Kerminen et al.^80^. At each merging step, we performed a permutation test by randomly reassigning individual copying vectors across the two groups being merged 1,000 times, preserving original group sizes. The empirical p-value was computed as the proportion of permuted TVD values greater than or equal to the observed TVD. Group pairs with p ≥ 0.01 were considered not significantly differentiated and collapsed into a single group before proceeding to the next merging step.

### EM-GMM

To assess the probability of each ancient individual belonging to either cluster 1 or cluster 2, we applied a semi-supervised Gaussian mixture model (GMM) on the Chromopainter coancestry matrix (chunklengths.out), following the approach described in Kerminen et al.^80^. Chunk lengths rather than chunk counts were used to mitigate the potential impact of phase switch errors in ancient individual phasing. Chromopainter was run using as recipients, ancient individuals assigned to Leiden clusters ‘1_1_1_1’ or ‘1_1_2_1’, as well as present-day individuals assigned to cluster 1 or cluster 2 in the FS*_south central_* analysis, and using as donors only present-day individuals from cluster 1 and cluster 2. For each ancient individual i, we calculated the log-ratio of average chunk length copied from cluster 1 versus cluster 2 reference individuals:

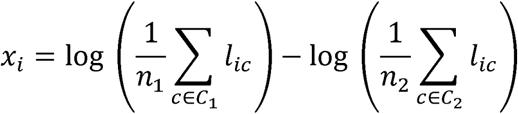

where n_1_ and n_2_ are the number of present-day reference individuals assigned to cluster 1 and cluster 2 respectively, and l_ic_ is the total chunk length that individual i copied from reference individual c. A supervised GMM was then fitted on the x_i_ values using an EM algorithm to assign each ancient individual a posterior probability of belonging to either cluster.

### SOURCEFINDv2

SOURCEFIND^57^ identifies the reference clusters with which each target individual shares the most recent ancestry and estimates their relative proportions using MCMC, while accounting for potential biases in the Chromopainter analysis, such as differences in sample sizes among potential source groups. To implement this, we first ran CHROMOPAINTER in donor mode (-f mode), re-estimated mutation probability and switch rate parameters following the same procedure described above (chromosomes 1, 8, 15, and 22), before fixing these parameters for the final painting run. SOURCEFIND models the number of source groups contributing ancestry to target individuals using a truncated Poisson prior. We set the mean of this prior to 4, while allowing up to 8 source groups to contribute in each MCMC iteration, using default parameters for all other settings. The first 50,000 MCMC iterations out of 200,000 were discarded as burn-in. Thereafter, we sampled mixture coefficients every 1,000 iterations and computed the final ancestry proportions by averaging these coefficients across the posterior samples.

### Random forest

To formally quantify the separation between cluster 1 and cluster 2, we trained a random forest classifier using per-individual SOURCEFIND ancestry proportions as predictors, implemented with the randomForest R package^81^. Near-zero variance predictors were removed prior to training. The dataset was split into a training set (80%) and a held-out test set (20%) using stratified sampling. The number of variables sampled at each split (mtry) was tuned by selecting the value minimising out-of-bag error across a grid of candidate values of number of predictors. The final model was trained with 1,000 trees using the best mtry value. Model performance was evaluated on the held-out test set and additionally assessed using leave-one-out cross-validation on the full dataset. Variable importance was quantified using mean decrease in Gini impurity and mean decrease in accuracy.

### FastGLOBETROTTER

Admixture dates were estimated using fastGLOBETROTTER^58^. For this analysis, ChromoPainter was run in donor mode, with south central African populations defined by FS*_south central_* as recipients (south central groups were merged based on the TVD permutation test described above) and all populations used in the second SOURCEFIND run as both recipients and donors. For each target group, 100 bootstrap replicates were performed to obtain 95% confidence intervals for admixture time. The “Null.ind” option was set to 1.

### IBD

IBD segments between pairs of present-day individuals were detected with hap-IBD^82^ using default parameters. Breaks and gaps in IBD segments were removed using the merge-ibd-segments program^83^, merging segments that overlapped or were separated by a gap ≤ 0.6 cM and contained at most one discordant homozygous genotype. Analyses were restricted to a set of predefined genomic regions excluding telomeres and centromeres^84^. One region on chromosome 19 was excluded after showing a significant depletion of IBD segments relative to expectation based on region length. We then generated a matrix of pairwise cumulative IBD length between individuals, considering only IBD segments ≥ 4 cM.

Population-level cluster labels were defined as described above, using Leiden cluster assignments for external populations and FS*_south central_* assignments for south central African individuals. Mean cumulative length of IBD (with IBD segments ≥ 4 cM) was computed across all possible pairs of individuals (including pairs sharing zero IBD) between population-cluster groups, then averaged within each geographic region (Angola/Namibia, southern Bantu, and Great Lakes Bantu). For DRC populations, comparisons were kept at the population-cluster level without regional averaging.

To assess uncertainty, we performed a block bootstrap over genomic regions (n = 1,000 iterations). At each iteration, regions were resampled with replacement and cumulative IBD lengths recomputed accordingly. Bootstrap 95% confidence intervals were derived from the 2.5th and 97.5th percentiles of the bootstrap distribution. P-values were derived from the bootstrap distribution of differences between groups. One-sided tests were used for the three serial founder model tests given their a priori directional predictions; two-sided tests were used elsewhere. All p-values were adjusted for multiple testing using the Benjamini-Hochberg procedure.

## Acknowledgments

We thank Alison Barton, Romain Fournier, Barbara Teixeira de Sousa Coelho Mota and Esther Brielle for helpful discussions on various aspects of this work. Douglas Kennett, Brendan Culleton, and Laurie Eccles for radiocarbon dating support. Nicole Adamski, Nasreen Broomandkhoshbacht, Elizabeth Curtis, Matthew Ferry, Kristin Stewardson and Fatma Zalzala for wet lab support. Rebecca Bernardos for sampling handing. Inigo Olalde and Iosif Lazaridis for bioinformatics support. We thank the participants of different dataset used in this study as well as the Livingstone Museum and National Heritage Conservation Commission in Zambia and the Malawi Department of Museums & Monuments (MDMM). We acknowledge the Research Computing Group at Harvard Medical School for their support. The aDNA analysis was funded by NIH grant HG012287; John Templeton Foundation grant 61220; a gift from Jean-Francois Clin; the Allen Discovery Center program, a Paul G. Allen Frontiers Group advised program of the Allen Family Philanthropies; and the Howard Hughes Medical Institute (HHMI).

## References

1. Derricourt, R.M. (1976). The Chronology of Zambian Prehistory. Transafrican J. Hist. 5, 1–31.

2. de Luna, K.M. (2012). Surveying the Boundaries of Historical Linguistics and Archaeology: Early Settlement in South Central Africa. Afr. Archaeol. Rev. 29, 209–251. 10.1007/s10437-012-9112-1.

3. Musonda, F.B. (1984). Late Pleistocene and Holocene Microlithic Industries from the Lunsemfwa Basin, Zambia. South Afr. Archaeol. Bull. 39, 24–36. 10.2307/3888592.

4. Juwayeyi, Y. (2020). Archaeology and Oral Tradition in Malawi: Origins and Early History of the Chewa (UCT Press).

5. Katongo, M., Fleisher, J.B., and Prendergast, M.E. (2025). Hunting, Fishing, and Herding in Later Stone Age and Iron Age Zambia: A Review of Zooarchaeological Evidence. Afr. Archaeol. Rev. 42, 143–172. 10.1007/s10437-025-09612-0.

6. Phillipson, D.W. (1976). The prehistory of Eastern Zambia (British institue in Eastern Africa).

7. Barham, L., and Mitchell, P. (2008). The first Africans: African archaeology from the earliest tool makers to most recent foragers (Cambridge University Press).

8. Grollemund, R., Branford, S., Bostoen, K., Meade, A., Venditti, C., and Pagel, M. (2015). Bantu expansion shows that habitat alters the route and pace of human dispersals. Proc. Natl. Acad. Sci. 112, 13296–13301. 10.1073/pnas.1503793112.

9. De Luna, K.M. (2017). Collecting food, cultivating people: subsistence and society in Central Africa (Yale University Press).

10. Grollemund, R., Schoenbrun, D., and Vansina, J. (2023). Moving Histories: Bantu Language Expansions, Eclectic Economies, and Mobilities. J. Afr. Hist. 64, 13–37. 10.1017/S0021853722000780.

11. Bostoen, K., Coutros, P., Doman, J., and Matonda Sakala, I.R. (2025). Rethinking the Bantu expansion from south of the Congo rainforest. In An archaeology of the Bantu expansion : early settlers south of the Congo rainforest (Routledge), pp. 565–584. 10.4324/9781032658148-29.

12. Fagan, B.M. (1969). Early Trade and Raw Materials in South Central Africa. J. Afr. Hist. 10, 1–13.

13. Stephens, J., Renson, V., Killick, D., Kaliba, P., and Matundu, S. (2026). Long distance trade of copper ingots from the eastern Copperbelt (Democratic Republic of Congo) to central Malawi. J. Archaeol. Sci. Rep. 73, 105845. 10.1016/j.jasrep.2026.105845.

14. Schadeberg, T.C. (1994). Spirantization and the 7-to-5 Vowel Merger in Bantu. Belg. J. Linguist. 9, 73–84. 10.1075/bjl.9.06sch.

15. de Maret, P. (1979). Luba Roots: The First Complete Iron Age Sequence in Zaϊre. Curr. Anthropol. 20, 233–235.

16. Macola, G. (2015). Luba–Lunda states. In The Encyclopedia of Empire, N. Dalziel and J. M. MacKenzie, eds. (Wiley), pp. 1–6. 10.1002/9781118455074.wbeoe060.

17. Vansina, J. (1975). Kingdoms of the Savanna a history of Central African states until European occupation 1966 Repr. (Univ. of Wisconsin Press).

18. Thornton, J.K. (2020). A history of West Central Africa to 1850 (Cambridge University Press).

19. Schoffeleers, J.M. (1992). River of blood: the genesis of a martyr cult in Southern Malawi, c. A.D. 1600 (Univ. of Wisconsin Press).

20. Choudhury, A., Sengupta, D., Ramsay, M., and Schlebusch, C. (2021). Bantu-speaker migration and admixture in southern Africa. Hum. Mol. Genet. 30, R56–R63. 10.1093/hmg/ddaa274.

21. Tallman, S., Sungo, M. das D., Saranga, S., and Beleza, S. (2023). Whole genomes from Angola and Mozambique inform about the origins and dispersals of major African migrations. Nat. Commun. 14, 7967. 10.1038/s41467-023-43717-x.

22. Fortes-Lima, C.A., Burgarella, C., Hammarén, R., Eriksson, A., Vicente, M., Jolly, C., Semo, A., Gunnink, H., Pacchiarotti, S., Mundeke, L., et al. (2024). The genetic legacy of the expansion of Bantu-speaking peoples in Africa. Nature 625, 540–547. 10.1038/s41586-023-06770-6.

23. Breton, G., Barham, L., Mudenda, G., Soodyall, H., Schlebusch, C.M., and Jakobsson, M. (2024). BaTwa populations from Zambia retain ancestry of past hunter-gatherer groups. Nat. Commun. 15, 7307. 10.1038/s41467-024-50733-y.

24. Morris, A.G., and Ribot, I. (2006). Morphometric cranial identity of prehistoric Malawians in the light of sub-Saharan African diversity. Am. J. Phys. Anthropol. 130, 10–25. 10.1002/ajpa.20308.

25. Morris, A.G. (2003). The Myth of the East African “Bushmen.” South Afr. Archaeol. Bull. 58, 85–90. 10.2307/3889305.

26. Morris, A.G. (2002). Isolation and the origin of the khoisan: Late pleistocene and early holocene human evolution at the southern end of Africa. Hum. Evol. 17, 231–240. 10.1007/BF02436374.

27. Hogg, A.G., Heaton, T.J., Hua, Q., Palmer, J.G., Turney, C.S., Southon, J., Bayliss, A., Blackwell, P.G., Boswijk, G., Ramsey, C.B., et al. (2020). SHCal20 Southern Hemisphere Calibration, 0–55,000 Years cal BP. Radiocarbon 62, 759–778. 10.1017/RDC.2020.59.

28. Ramsey, C.B. (2009). Bayesian Analysis of Radiocarbon Dates. Radiocarbon 51, 337–360. 10.1017/S0033822200033865.

29. Lipson, M., Sawchuk, E.A., Thompson, J.C., Oppenheimer, J., Tryon, C.A., Ranhorn, K.L., de Luna, K.M., Sirak, K.A., Olalde, I., Ambrose, S.H., et al. (2022). Ancient DNA and deep population structure in sub-Saharan African foragers. Nature 603, 290–296. 10.1038/s41586-022-04430-9.

30. Lipson, M., Ribot, I., Mallick, S., Rohland, N., Olalde, I., Adamski, N., Broomandkhoshbacht, N., Lawson, A.M., López, S., Oppenheimer, J., et al. (2020). Ancient West African foragers in the context of African population history. Nature 577, 665–670. 10.1038/s41586-020-1929-1.

31. Llorente, M.G., Jones, E.R., Eriksson, A., Siska, V., Arthur, K.W., Arthur, J.W., Curtis, M.C., Stock, J.T., Coltorti, M., Pieruccini, P., et al. (2015). Ancient Ethiopian genome reveals extensive Eurasian admixture in Eastern Africa. Science 350, 820–822. 10.1126/science.aad2879.

32. Wang, K., Goldstein, S., Bleasdale, M., Clist, B., Bostoen, K., Bakwa-Lufu, P., Buck, L.T., Crowther, A., Dème, A., McIntosh, R.J., et al. (2020). Ancient genomes reveal complex patterns of population movement, interaction, and replacement in sub-Saharan Africa. Sci. Adv. 6, eaaz0183. 10.1126/sciadv.aaz0183.

33. Prendergast, M.E., Lipson, M., Sawchuk, E.A., Olalde, I., Ogola, C.A., Rohland, N., Sirak, K.A., Adamski, N., Bernardos, R., Broomandkhoshbacht, N., et al. (2019). Ancient DNA reveals a multistep spread of the first herders into sub-Saharan Africa. Science 365, eaaw6275. 10.1126/science.aaw6275.

34. Schlebusch, C.M., Malmström, H., Günther, T., Sjödin, P., Coutinho, A., Edlund, H., Munters, A.R., Vicente, M., Steyn, M., Soodyall, H., et al. (2017). Southern African ancient genomes estimate modern human divergence to 350,000 to 260,000 years ago. Science 358, 652–655. 10.1126/science.aao6266.

35. Skoglund, P., Thompson, J.C., Prendergast, M.E., Mittnik, A., Sirak, K., Hajdinjak, M., Salie, T., Rohland, N., Mallick, S., Peltzer, A., et al. (2017). Reconstructing Prehistoric African Population Structure. Cell 171, 59–71.e21. 10.1016/j.cell.2017.08.049.

36. Fan, S., Kelly, D.E., Beltrame, M.H., Hansen, M.E.B., Mallick, S., Ranciaro, A., Hirbo, J., Thompson, S., Beggs, W., Nyambo, T., et al. (2019). African evolutionary history inferred from whole genome sequence data of 44 indigenous African populations. Genome Biol. 20, 82. 10.1186/s13059-019-1679-2.

37. Bergström, A., McCarthy, S.A., Hui, R., Almarri, M.A., Ayub, Q., Danecek, P., Chen, Y., Felkel, S., Hallast, P., Kamm, J., et al. (2020). Insights into human genetic variation and population history from 929 diverse genomes. Science 367, eaay5012. 10.1126/science.aay5012.

38. Mallick, S., Li, H., Lipson, M., Mathieson, I., Gymrek, M., Racimo, F., Zhao, M., Chennagiri, N., Nordenfelt, S., Tandon, A., et al. (2016). The Simons Genome Diversity Project: 300 genomes from 142 diverse populations. Nature 538, 201–206. 10.1038/nature18964.

39. Meyer, M., Kircher, M., Gansauge, M.-T., Li, H., Racimo, F., Mallick, S., Schraiber, J.G., Jay, F., Prüfer, K., de Filippo, C., et al. (2012). A High-Coverage Genome Sequence from an Archaic Denisovan Individual. Science 338, 222–226. 10.1126/science.1224344.

40. Tishkoff, S.A., Reed, F.A., Friedlaender, F.R., Ehret, C., Ranciaro, A., Froment, A., Hirbo, J.B., Awomoyi, A.A., Bodo, J.-M., Doumbo, O., et al. (2009). The Genetic Structure and History of Africans and African Americans. Science 324, 1035–1044. 10.1126/science.1172257.

41. Bird, N., Ormond, L., Awah, P., Caldwell, E.F., Connell, B., Elamin, M., Fadlelmola, F.M., Matthew Fomine, F.L., López, S., MacEachern, S., et al. (2023). Dense sampling of ethnic groups within African countries reveals fine-scale genetic structure and extensive historical admixture. Sci. Adv. 9, eabq2616. 10.1126/sciadv.abq2616.

42. Patin, E., Siddle, K.J., Laval, G., Quach, H., Harmant, C., Becker, N., Froment, A., Régnault, B., Lemée, L., Gravel, S., et al. (2014). The impact of agricultural emergence on the genetic history of African rainforest hunter-gatherers and agriculturalists. Nat. Commun. 5, 3163. 10.1038/ncomms4163.

43. Choudhury, A., Aron, S., Botigué, L.R., Sengupta, D., Botha, G., Bensellak, T., Wells, G., Kumuthini, J., Shriner, D., Fakim, Y.J., et al. (2020). High-depth African genomes inform human migration and health. Nature 586, 741–748. 10.1038/s41586-020-2859-7.

44. Derricourt, R.M. (1985). Man on the Kafue: the archaeology and history of the Itezhitezhi area of Zambia (Ethnographica).

45. Haak, W., Lazaridis, I., Patterson, N., Rohland, N., Mallick, S., Llamas, B., Brandt, G., Nordenfelt, S., Harney, E., Stewardson, K., et al. (2015). Massive migration from the steppe was a source for Indo-European languages in Europe. Nature 522, 207–211. 10.1038/nature14317.

46. Patterson, N., Moorjani, P., Luo, Y., Mallick, S., Rohland, N., Zhan, Y., Genschoreck, T., Webster, T., and Reich, D. (2012). Ancient Admixture in Human History. Genetics 192, 1065–1093. 10.1534/genetics.112.145037.

47. Harney, É., Patterson, N., Reich, D., and Wakeley, J. (2021). Assessing the performance of qpAdm: a statistical tool for studying population admixture. Genetics 217, iyaa045. 10.1093/genetics/iyaa045.

48. Chintalapati, M., Patterson, N., and Moorjani, P. (2022). The spatiotemporal patterns of major human admixture events during the European Holocene. eLife 11, e77625. 10.7554/eLife.77625.

49. Ehret, C. (2001). An African classical age: eastern and southern Africa in world history, 1000 B.C. to A.D. 400 1st pbk. ed. (University Press of Virginia ; J. Currey).

50. Pawlowicz, M., Fleisher, J., and de Luna, K. (2020). Capturing People on the Move: Spatial Analysis and Remote Sensing in the Bantu Mobility Project, Basanga, Zambia. Afr. Archaeol. Rev. 37, 69–93. 10.1007/s10437-020-09363-0.

51. Traag, V.A., Waltman, L., and van Eck, N.J. (2019). From Louvain to Leiden: guaranteeing well-connected communities. Sci. Rep. 9, 5233. 10.1038/s41598-019-41695-z.

52. Lawson, D.J., Hellenthal, G., Myers, S., and Falush, D. (2012). Inference of Population Structure using Dense Haplotype Data. PLoS Genet. 8, e1002453. 10.1371/journal.pgen.1002453.

53. Leslie, S., Winney, B., Hellenthal, G., Davison, D., Boumertit, A., Day, T., Hutnik, K., Royrvik, E.C., Cunliffe, B., Lawson, D.J., et al. (2015). The fine-scale genetic structure of the British population. Nature 519, 309–314. 10.1038/nature14230.

54. Genetic legacy of state centralization in the Kuba Kingdom of the Democratic Republic of the Congo | PNAS https://www.pnas.org/doi/10.1073/pnas.1811211115.

55. Brielle, E.S., Fleisher, J., Wynne-Jones, S., Sirak, K., Broomandkhoshbacht, N., Callan, K., Curtis, E., Iliev, L., Lawson, A.M., Oppenheimer, J., et al. (2023). Entwined African and Asian genetic roots of medieval peoples of the Swahili coast. Nature 615, 866–873. 10.1038/s41586-023-05754-w.

56. Bostoen, K. (2025). Language and history in Bantu-speaking Africa. In The Oxford Guide to the Bantu Languages, L. Marten, E. Hurst-Harosh, N. C. Kula, and J. Zeller, eds. (Oxford University PressOxford), pp. 17–27. 10.1093/oso/9780198808343.003.0003.

57. Chacón-Duque, J.-C., Adhikari, K., Fuentes-Guajardo, M., Mendoza-Revilla, J., Acuña-Alonzo, V., Barquera, R., Quinto-Sánchez, M., Gómez-Valdés, J., Everardo Martínez, P., Villamil-Ramírez, H., et al. (2018). Latin Americans show wide-spread Converso ancestry and imprint of local Native ancestry on physical appearance. Nat. Commun. 9, 5388. 10.1038/s41467-018-07748-z.

58. Hellenthal, G., Busby, G.B.J., Band, G., Wilson, J.F., Capelli, C., Falush, D., and Myers, S. (2014). A Genetic Atlas of Human Admixture History. Science 343, 747–751. 10.1126/science.1243518.

59. Goldstein, S.T., Crowther, A., Henry, E.R., Janzen, A., Katongo, M., Brown, S., Farr, J., Le Moyne, C., Picin, A., Richter, K.K., et al. (2021). Revisiting Kalundu Mound, Zambia: Implications for the Timing of Social and Subsistence Transitions in Iron Age Southern Africa. Afr. Archaeol. Rev. 38, 625–655. 10.1007/s10437-021-09440-y.

60. Katanekwa, N.M. (2016). The prehistory of the 73+ Bantu languages and Bantu language groups of Zambia 3000 BC to 1600 AD: (including a new interpretation of the classification, origins and migrations of the 600+ Bantu languages and Bantu language groups of Africa (Lendekwa Heritage Consultancy & Services).

61. Gibbon, V.E., Gallagher, A., and Huffman, T.N. (2014). Bioarchaeological Analysis of Iron Age Human Skeletons from Zambia. Int. J. Osteoarchaeol. 24, 100–110. 10.1002/oa.2231.

62. McKeeby, Z. (2024). Mapping the Iron Age in Southern Africa: Magnetometry at two Iron Age villages in Western Zambia. J. Archaeol. Sci. 163, 105937. 10.1016/j.jas.2024.105937.

63. Kingdoms of South-Central Africa: Sources, Historiography, and History | Oxford Research Encyclopedia of African History | Oxford Academic https://academic.oup.com/edited-volume/61663/chapter-abstract/553478632?redirectedFrom=fulltext.

64. Gordon, D.M. (2023). The Quotidian Politics of a Love Story: Researching, Assembling, and Mobilizing the Lunda Legend in the Late Nineteenth Century. J. Afr. Hist. 64, 209–228. 10.1017/S0021853723000300.

65. Mainga, M. (1973). Bulozi under the Luyana kings: political evolution and state formation in pre-colonial Zambia (Longmans).

66. Pikirayi, I. (1993). The archaeological identity of the Mutapa State: towards an historical archaeology of northern Zimbabwe (Societas Archaeologica Upsaliensis□: Distributedby Dept. of Archaeology, Uppsala University).

67. Fortune, G., and Kashoki, M.E. (2001). An outline of Silozi grammar New ed. (Bookworld Publishres).

68. Van (2016). Tears Of Rain - Ethnicity & Hist (Taylor and Francis).

69. Vansina, J. (2004). How societies are born: governance in West Central Africa before 1600 (University of Virginia Press).

70. Ramsey, C.B. (2009). Bayesian Analysis of Radiocarbon Dates. Radiocarbon 51, 337–360. 10.1017/S0033822200033865.

71. Hogg, A.G., Heaton, T.J., Hua, Q., Palmer, J.G., Turney, C.S., Southon, J., Bayliss, A., Blackwell, P.G., Boswijk, G., Ramsey, C.B., et al. (2020). SHCal20 Southern Hemisphere Calibration, 0–55,000 Years cal BP. Radiocarbon 62, 759–778. 10.1017/RDC.2020.59.

72. Rohland, N., Mallick, S., Mah, M., Maier, R., Patterson, N., and Reich, D. (2022). Three assays for in-solution enrichment of ancient human DNA at more than a million SNPs. Genome Res. 32, 2068–2078. 10.1101/gr.276728.122.

73. Fournier, R., Fulton, A.P., and Reich, D. (2025). A SNP panel for co-analysis of capture and shotgun ancient DNA data. Preprint at bioRxiv, 10.1101/2025.07.30.667733 https://doi.org/10.1101/2025.07.30.667733.

74. Manichaikul, A., Mychaleckyj, J.C., Rich, S.S., Daly, K., Sale, M., and Chen, W.-M. (2010). Robust relationship inference in genome-wide association studies. Bioinformatics 26, 2867–2873. 10.1093/bioinformatics/btq559.

75. Rubinacci, S., Ribeiro, D.M., Hofmeister, R.J., and Delaneau, O. (2021). Efficient phasing and imputation of low-coverage sequencing data using large reference panels. Nat. Genet. 53, 120–126. 10.1038/s41588-020-00756-0.

76. Akbari, A., Perry, A., Barton, A.R., Kariminejad, M., Gazal, S., Li, Z., Zeng, Y., Mittnik, A., Patterson, N., Mah, M., et al. (2026). Ancient DNA reveals pervasive directional selection across West Eurasia. Nature 654, 419–428. 10.1038/s41586-026-10358-1.

78. Patterson, N., Price, A.L., and Reich, D. (2006). Population Structure and Eigenanalysis. PLoS Genet. 2, e190. 10.1371/journal.pgen.0020190.

79. Csárdi, G., and Nepusz, T. (2006). The igraph software package for complex network research. In.

80. Kerminen, S., Havulinna, A.S., Hellenthal, G., Martin, A.R., Sarin, A.-P., Perola, M., Palotie, A., Salomaa, V., Daly, M.J., Ripatti, S., et al. (2017). Fine-Scale Genetic Structure in Finland. G3 GenesGenomesGenetics 7, 3459–3468. 10.1534/g3.117.300217.

81. Liaw, A., and Wiener, M. (2007). Classification and Regression by randomForest. In.

82. Zhou, Y., Browning, S.R., and Browning, B.L. (2020). A Fast and Simple Method for Detecting Identity-by-Descent Segments in Large-Scale Data. Am. J. Hum. Genet. 106, 426–437. 10.1016/j.ajhg.2020.02.010.

83. Browning, B.L., and Browning, S.R. (2013). Improving the Accuracy and Efficiency of Identity-by-Descent Detection in Population Data. Genetics 194, 459–471. 10.1534/genetics.113.150029.

84. Nait Saada, J., Kalantzis, G., Shyr, D., Cooper, F., Robinson, M., Gusev, A., and Palamara, P.F. (2020). Identity-by-descent detection across 487,409 British samples reveals fine scale population structure and ultra-rare variant associations. Nat. Commun. 11, 6130. 10.1038/s41467-020-19588-x.

