## Supplementary Material for "The south Congo Basin was critical to Bantu settlement of south central Africa"

**Permissions and sampling protocols**

The Livingstone Museum (LM) in Zambia is the repository for all sites from Zambia included in this study, plus all but one sample from the site of Nkhudzi Bay in Malawi. M.E.P. and E.A.S. received permissions to sample and to export skeletal remains for destructive sampling from the Director of the LM and from the National Heritage Conservation Commission (Permit NHCC8WR/004/17). All remaining skeletal tissue samples from the LM were repatriated to that institution in June 2018 and June 2019. Remaining powder, DNA extracts, and libraries remain under curation at the Reich Laboratory at Harvard University.

At the LM, skeletal remains were inventoried by A.M., M.E.P., and E.A.S. prior to sampling. For some human remains, little contextual information was available beyond a site name, the name of a landowner or donor, and/or a rough location. The museum supported analysis of these remains in order to improve information about poorly documented collections. Sampling for ancient DNA and radiocarbon dating was aimed first, at well-published Iron Age (IA) archaeological sites; and second, at sampling underdocumented remains. We avoided sampling Later Stone Age (LSA) to IA sites where there was prior sampling or active plans to sample for aDNA by other researchers: this includes Gwisho Hot-Springs, Makwe Rockshelter, Mumbwa Caves, Nachikufu Rockshelter, and Thandwe Rockshelter.

Many collections were fragmentary, heavily commingled, and had minimal or conflicting labels. We aimed to sample unique individuals, based on morphology, though eight samples were later shown to be duplicates representing four individuals. One or two petrous, tooth, or limb bone samples were collected per individual (denoted .01, .02); often the second sample was collected to enable radiocarbon dating in case of working aDNA, and in many cases was left unsampled. Samples with an antimere (opposite side pair), or that were already fragmentary, were preferentially chosen to minimize impacts. We deposited digital and photographic databases with the museum that document our skeletal inventories and sampling activity.

**Archaeological Site Summaries**

Here we provide summaries for the archaeological sites and burial contexts for newly reported individuals, including those for whom genetic data was sequenced but excluded from our analysis (**Supplementary Table 1**). Latitudes and longitudes are sometimes approximate, and additional information may be obtained from relevant museums or cultural heritage authorities. Enhanced libraries are reported here from individuals whose genomes were previously published at Fingira^1^ and Hora^2^; details are provided in those publications. We do not include here descriptions of sites for which there were no successful aDNA results: these include Leopard’s Hill Cave, Luangwa Valley Game Reserve, Nsalu Caves (Mpika), Sanjika Hill, and Sinachilundu. Dates are presented in site descriptions as calibrated years before/in the common era (cal BCE/CE), and raw dates and further details can be found in **Supplementary Table 2** and in the radiocarbon dating section below.

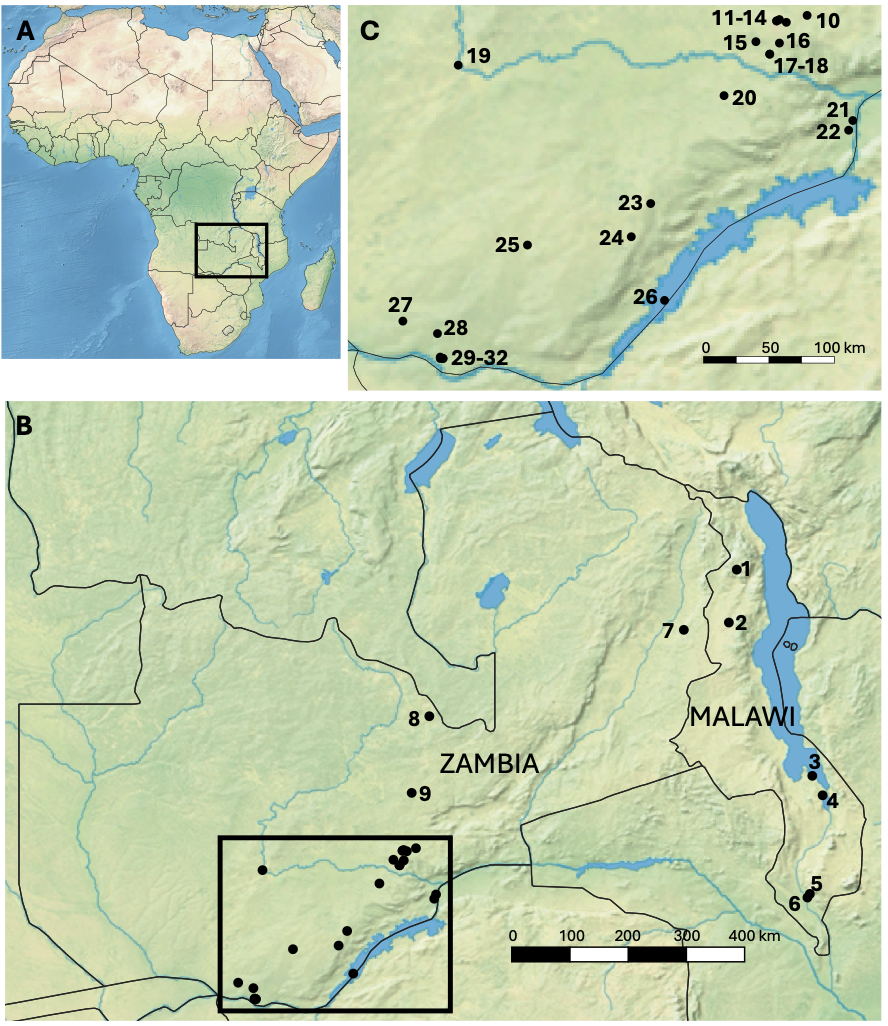

**Figure S1.** Map of Africa (**A**) showing study area in Zambia and Malawi (**B**), with detail of the Lusaka and Livingstone areas in **C**. 1, Fingira; 2, Hora; 3, Nkudzi Bay; 4, Mtemankhokwe; 5, Dzimbiri; 6, Chipakusa; 7, Kanyankunde; 8, Fiwale Mission; 9, Moresby-White Farm; 10, Silver Rest Farm; 11, Olympia Park; 12, Mayburgh's Farm; 13, Mr. Bartlett's Garden; 14, Lusaka Twin Palm Road; 15, Twin Rivers; 16, Freeman's (Kapongo) Cave; 17, Chipongwe; 18, Shimabala; 19, Kalala Island; 20, Kelly's Farm; 21, Lusitu/Zimbe; 22, Ingombe Ilede; 23, Ndonde; 24, Sikalongo; 25, Kalomo; 26, Sinachisingili; 27, Simbusenga; 28, Sinde Mission; 29, Batoka Round-about; 30, Linda Compound; 31, Nansanzu School; 32, Water Tower.

**Batoka Round-about, Zambia**

*Southern Province, Latitude -17.84 Longitude 25.85*

*Primary reference: none*

This is a poorly documented site, presumed to be at the roundabout of the same name today in Livingstone (we assigned those coordinates). Human remains were limited to poorly preserved long bone shafts and fifteen isolated teeth, consistent with a single individual around 20 years of age. We sampled this individual via a tooth (BR.01.02), which produced working aDNA (I10840), confirmed male genetic sex, and was directly dated to the 18th-20th centuries CE.

**Chipakusa, Malawi**

*Thyolo District, Southern Region, Latitude -16.18 Longitude 34.93*

*Primary reference: none*

This site, also spelled Chipakuza, produced a large number of glass beads, dated via typology to c. 1700-1900 CE^3^. Two individuals, represented by a tooth and a petrous bone, respectively, were sampled from this site and both produced working aDNA (I6162, I6163). While no direct dates are available, glass beads give an indication of the site’s chronology.

**Chipongwe, Zambia**

*Lusaka Province, Latitude -15.65 Longitude 28.23.*

*Primary reference: Clark & Toerien 1955 [ref* ^4^*].*

Chipongwe is a sinkhole in a series of formations known as the Chipongwe Caves located near the Shimabala quarry, about 3.2 km west of the rail line, south of Lusaka. Human remains from the earliest expedition are curated in South Africa^5^. Additional remains were recovered in1952-53^4^, and some were sent to South Africa while others are at the LM.

Clark & Toerien (1955) noted substantial disturbance at the site. They identified remains of at least four individuals, likely representing shallow burials that were later disturbed and scattered downslope. Lithic technology was attributed to the LSA^6^. Faunal remains were dominated by wild bovids and suids, and pottery resembled recent local traditions. None of these materials could be confidently associated with the human remains. Clark & Toerien (1955) noted that deep caves were frequently used for refuge during late 19th century raids, and suggested the human remains might date to this time; they also suggested that the individuals buried there might have been the makers of LSA lithics, and argued the individuals may have been foragers unrelated to neighboring Bantu speakers.

The human remains curated in South Africa have been well described^4,7^, and one of these individuals recently formed part of a genetic study, producing a direct AMS date of 97±29 bp (Ua-61975, calibrated to 1698-1950 cal CE)^8^. This range is consistent with Clark & Toerien’s (1955) chronological interpretation, and contradicts de Villiers and Fatti’s (1990)[ref ^9^] attribution to c. 4000 BP.

We did not find remains in the LM that matched descriptions by Clark & Toerien (1955). However, at least two additional sets of human remains were deposited at the Livingstone by J.D. Clark (accession numbers 6986/6987). We found these remains heavily commingled, including with faunal remains, and with few labeled skeletal elements. If accession numbers 6986 and 6987 are separate excavation contexts, the MNI for Chipongwe at Livingstone could be seven individuals; more conservatively, there are at least six individuals represented across the two accessions: four adults represented by crania, an infant represented by cranial remains, and an older child represented by postcrania.

We collected eight samples from the seven individuals we had identified via morphology. Five of these produced ancient DNA, but these in fact only represent three unique individuals, reflecting extensive commingling.  The first individual (I10843, sample CH.02.01) is an adult male represented by a lower left central incisor in a mandibular fragment. The second individual is another adult male represented by an isolated lower left canine (I10842, CH.01.01) and a lower left P3 in a mandible with “Chipongwe 3” written on it (CH.04.01). The final individual (I8386) is an adult female represented by an intact skull except for the face labeled “IV cranium”, in which the petrous was sampled using cranial base drilling method (CH.07.01). An isolated worn lower left molar (CH.03.01) was found to be genetically the same individual.

Calibrated direct radiocarbon dates on these remains range from the 17th-20th centuries, with median dates falling in the late 17th to early 18th centuries. These overlap with but are generally earlier than the date produced by Fortes-Lima et al. (2024), suggesting repeated use of the site as a burial ground during the modern era, and possibly earlier than the late 19th century as originally suggested by Clark & Toerien (1955).

**Dzimbiri, Malawi**

*Thyolo District, Southern Region, Latitude -16.12 Longitude 34.97*

*Primary reference: none*

This site, also spelled Drimbiri, is located in the Lower Shire Valley, and produced a large number of glass beads, dated via typology to c. 1700-1900 CE^3^. Five individuals, represented by three teeth and two petrous bone samples, were sampled from this site. While the petrous bone samples failed, all three teeth produced working aDNA. Two of these belong to the same individual (I5777), and this and a second individual (I5776) are both genetically female. While no direct dates are available, glass beads give an indication of the site’s chronology.

**Fiwale Mission, Zambia**

*Copperbelt Province, Latitude -13.20 Longitude 28.72.*

*Primary reference: none.*

This site is located in Ndola District in Copperbelt Province, not to be confused with Fiwila Mission in Central Province. Two badly eroded skulls were present in the LM, denoted Fiwale 1 and 2. Fiwale 1 is a small adult skull with extensive taphonomic damage. We sampled the *in situ* petrous bone (FM.01.01), which produced working aDNA (I10723) from a genetic male. This individual was directly dated to the late 17th to early 19th centuries CE. The other skull (Fiwale 2) is complete, and had a label reading “skull with no info, must be Fiwali mission,” which was later changed to read “Fawila.” Given these ambiguities, we chose not to sample this skull.

**Freeman’s Cave II (Kapongo Cave), Zambia**

*Lusaka Province, Latitude -15.57 Longitude 28.30.*

*Primary reference: none.*

This site, now known as Kapongo Cave, is located in the Kafue Gorge south of Lusaka, and is relatively well documented in terms of geology and paleontology^10^. Here we retain the colonial name of Freeman’s Cave for consistency with museum records. Little archaeological information is available for this site, but the accession register notes it as LSA and records that potsherds and human remains were deposited by Clark. No mention is made of the site by Clark (1950)[ref ^7^], although nearby caves (Chipongwe and Leopard’s Hill) are described. although nearby caves (Chipongwe and Leopard’s Hill) are described. We found human remains consisting of commingled cranial and postcranial fragments, consistent with representing a single adult female. A left petrous bone attached to the temporal yielded working aDNA (I10730), and was directly dated to the late 17th-18th centuries.

**Ingombe Ilede, Zambia**

*Southern Province, Latitude -16.20 Longitude 28.80.*

*Primary references: Chaplin 1962[ref* ^11^*]; Fagan et al. 1969[ref* ^12^*]; McIntosh & Fagan 2017[ref* ^13^*]*

Ingombe Ilede, located in the Middle Zambezi River, is one of the best-known Recent IA sites in Zambia. The site was discovered during water tank construction in 1960^11^, and eleven burials were recovered. Gold, glass beads, and copper ingots indicated that Ingombe Ilede lay at the center of important trading networks^12^.

These burials, which in some cases have elaborate grave goods (notably Burials 1, 2, 3, and 8), have been well documented by Fagan et al. (1969), who later excavated an additional 31 individuals (mainly infants and children) in a cemetery to the south of the previously documented burials. In total, 46 burials have been recorded, the bulk of which are curated in South Africa^14^. Fagan and Phillipson’s (1969) [ref ^15^] dating of the site, based on charcoal, placed the occupation in the 14th-15th centuries; they suggested this might be a second occupation of the site, and did not rule out an earlier late first/early second millennium occupation.

In the LM collections, cotton burial cloth fragments attached to copper bangles associated with Burials 3 and 8 were directly AMS dated by McIntosh & Fagan (2017). These newly generated dates shift the end date of the site’s occupation to the 15th-17th centuries, a time of active exchange networks between Ingombe Ilede and the Swahili coast, the development of the Mutapa state, and the establishment of Portuguese trading posts along the Zambezi^13^.

At the LM, we were able to locate and sample fragmentary remains from Burials 4 and 8, as well as a potential third individual represented by isolated teeth. Burial 9 was in a public display case, and we did not sample it. Murphy (1996:75) [ref ^16^] recorded more material at the LM than we were able to find, noting elements from Burials 1-4, 6, and 8.

For Burial 4, since no cranial material could be identified, we sampled an ulna fragment which failed to produce aDNA (II4.01). We also sampled an unlabeled incisor from a bag of commingled remains, which we assigned an arbitrary individual number (II.2017.01.01); this, too, failed to produce aDNA. For Burial 8, we sampled the *in situ* petrous via the cranial base drilling method (II8.02), and this produced working aDNA (I8383), and confirmed the genetic sex as male. We use the cloth date published by McIntosh & Fagan (2017) for this individual (370±30 uncal bp, Beta-415034). Burial 8 is one of the most important burials, with elaborate grave goods (including copper crosses, a gong, a hammerhead, pottery, and chicken bones) and adornment including gold, glass, and shell beads and copper bangles^12,13^.

We also identified human remains from a separate and relatively recent excavation, deposited by the National Heritage Conservation Commission (NHCC) at the Livingstone Museum with a listed date of 2013. We were unable to find records related to these excavations, but in the museum collections, we found isolated teeth and a right temporal fragment with attached petrous, consistent in size with an adult. We sampled the petrous (Il2013.01.01), which produced working aDNA (I8860) and genetic female sex. A direct date on this individual falls at the turn of the 19th-20th centuries, suggesting this burial is unrelated to the earlier activities at Ingombe Ilede.

**Kalala Island, Zambia**

*Southern Province, Latitude -15.73 Longitude 25.98*

*Primary reference: Derricourt 1985[ref* ^17^*]*

Kalala Island is the southernmost island within the Middle Kafue River, just before it reaches the Kafue Flats. It sits at an important ecological boundary today between a game reserve and grasslands occupied by pastoralists, and is near important IA occupations documented at mound sites in the Kafue Flats^18^.  In advance of dam construction, Derricourt (1985) documented more than 150 sites, many of them from the LSA. Excavations in 1974-75 focused on more recent deposits on Kalala Island, parts of which are submerged today. Excavations at Kalala Island Main site revealed the remains of at least 15 pole-and-daga huts, as well as hearths and ash lenses, thought to date to the Middle and Recent Iron Age, and overlying evidence of LSA and Early IA activities. Faunal remains indicated fishing and hunting, and early reports of cattle were later suggested to be buffalo^17^. Hippopotamus and elephant remains, along with ivory artifacts, suggest that ivory production was important.

Three radiocarbon dates on charcoal from Kalala Main Site were in reverse stratigraphic order but overlapping, suggesting a span of the mid 8th to late 9th centuries^17^. Based on ceramics, Derricourt suggested a chronology extending into the 10th century, while the presence of trade beads - some of which connect the site to the Swahili coast  – suggested additional later use. Eight burials were documented. Derricourt (1985:65) noted that though they appeared to be in undisturbed deposits, they were found with glass beads from the 17th century or later.

The burials are well described by de Villiers in Derricourt (1985). Two adult burials were poor in grave goods and were interpreted as hasty (KAL J and KAL EX; the latter also recorded as KAL E Burial 1). Others were lightly to extensively adorned, including an adult male with iron bands (KAL D), a young adult female with a dozen ivory armlets and a glass bead necklace (KAL I), another adult female with a necklace (KAL L Burial 1), and three infants with necklaces (KAL E Burial 2, KAL E Burial 3; and KAL L Burial 2).

In the LM, Kalala Island remains are curated together and could be largely connected to the individuals studied by de Villiers, even though commingled. We were unable to locate one individual (KAL L Burial 2), though we believe it may be commingled in a bag with KAL L Burial 1. We sampled the other seven individuals present, and all samples produced working aDNA. Of these, six had sufficient collagen for direct radiocarbon dates. These dates suggest a complex chronology with multiple burial events as early as the 14th century (KAL D) and 16th-17th century (KAL EX); the latter is the individual shown in the present study to be genetically distinctive, with strong affinities to forager groups (KAL EX; I8379). This person was buried in the purportedly most recent archaeological phase (Phase 9, Recent IA), with no grave goods or personal adornment, making it distinctive from the others. The other dated burials fall in the 18th-20th centuries. Collectively, none of the dated burials align with the stratigraphic phasing provided by Derricourt (1985), nor with the charcoal dates, confirming Derricourt’s view that the burials are intrusive and unrelated to earlier site use.

**Kalomo, Zambia**

*Southern Province, Latitude -17.03 Longitude 26.48*

*Primary reference: none*

While Kalomo is an archaeological culture of the IA, represented by a series of mounds there is no specific “Kalomo” burial mentioned in Fagan’s (Fagan 1967)[ref ^19^] work on this culture. Rather, the burial we studied appears to come from much earlier work.  An accession register reads only: *“Clark. Kalomo – 1 km north of. Iron Age. See ‘Stone Age Cultures…’”*. The coordinates provided here are thus imprecise, and we cannot identify an associated publication besides Clark (1950)[ref ^7^], who describes explorations of the geology and archaeology of the Kalomo River region. Clark (1950:148) also mentions two human skeletons recovered in association with pottery near the town of Kalomo: one by Gear (1926)[ref ^20^], which was later studied by Schepers (1935)[ref ^21^]; and another presumably distinct skeleton reported in a short footnote by Clark (1942:179)[ref ^22^]. The remains in LM most likely correspond to the Clark publication.

We sampled two individuals under the accession number 6980/SK5 for “Kalomo.” One is an adult male, from which we sampled a petrous bone (KM.01.01), which produced working aDNA (I8868), confirmed male sex, and dates to the 18th-20th centuries. A second individual is a juvenile represented by a tibia, and this sample (KM.02.01) produced no working aDNA.

Fortes-Lima et al. (2024)[ref ^8^] report aDNA data from an individual from “Horton's farm, Kalomo, Zambia” (museum accession number A4155). That individual is directly dated to 1635-1950 calCE (236±36 uncal bp, Ua-61830), which is consistent with the date we obtained on the Kalomo individual in the present study. We are uncertain whether these are the materials previously studied by Schepers (1935)[ref ^21^] or could also be from the Clark (1942)[ref ^22^] expedition. Horton was a large-scale landowner in the Choma-Kalomo area, so this does not offer much resolution.

**Kanyankunde Rockshelter, Zambia**

*Eastern Province, Latitude -11.78 Longitude 32.90*

*Primary reference: Macrae & Lancaster 1937[ref* ^23^*]; Phillipson 1976[ref* ^24^*]*

Kanyankunde Rockshelter is an LSA site located in the hills of the Luangwa Escarpment, to the northwest of a settlement at Lundazi. This site was briefly reported by Macrae & Lancaster (1937). Lancaster conducted a small excavation in 1936, recovering nondescript “bones”; the finds overall were likened to those studied by Dart and Del Grande (1931)[ref ^5^] at Mumbwa Cave. The shelter has a faint red painting, and deposits up to 90 cm thick separated into earlier and later phases. These included both LSA microlithic (Mode 5) technology and pottery, with differences noted in the earlier and later pottery deposits^24^. We identified cranial and postcranial remains consistent with a single adult female, with squatting facets on the tibia and antemortem incisor removal or loss. We sampled a petrous bone (KK.01.02) from this person, which yielded aDNA (I10738) and a direct date of 1425-1455 cal CE, which is notably both earlier and more tightly constrained than many other dates on individuals in the present study.

**Kelly’s Farm, Zambia**

*Southern Province, Latitude -15.95 Longitude 27.90*

*Primary reference: Fagan & Phillipson 1965[ref* ^25^*]*

This is a Recent IA site located near Mazabuka in Southern Province. Fagan & Phillipson (1965:270) provide a brief report of the site, interpreting it as a former Plateau Tonga village, likely abandoned in the early 20th century. They collected 61 potsherds, noting their similarities to modern Tonga pottery and suggesting they might be slightly later than those of Nabombe. No mention is made of human remains. At the Livingstone Museum, several long bones from this site were found, and their size and preservation is consistent with representing a single individual. We sampled an ulna (KF.01.01), which yielded aDNA (I10837) and confirmed this person as a genetic female. A direct date places this individual in the 18th-20th centuries.

**Linda Compound, Maramba, Zambia**

*Southern Province, Latitude -17.85 Longitude 25.87*

*Primary reference: None*

Accession information for this burial reads “*Linda Compound, Maramba, Livingstone, in house foundations, ?modern, assoc. 2 pots*.” This is not likely related to the Maramba Game Park Site, with a LSA burial documented by Clark (1950:112-113)[ref ^7^],  as those remains were heavily fossilized and were sent to South Africa for analysis. Linda is also an Early IA archaeological site documented by Vogel (1973) [ref ^26^], but this site description does not match either as it is a school, not a house, and remains were not found while digging foundations; additionally, Vogel mentions no burial. While other burials were also found in the Linda Compound area by Vogel (1971) (1971)[ref ^27^], these are clearly named Nansanzu School. It thus seems that this Linda Compound burial is archaeologically undocumented. Human remains are fragmented and consistent with one adult. We sampled a petrous bone (LC.01.01), generating working aDNA (I10729) from a genetic male, but there was insufficient collagen for a direct date.

**Lusaka Twin Palm Road, Zambia**

*Lusaka Province, Latitude -15.42 Longitude 28.35*

*Primary reference: None*

No archaeological information was available for this site, and we have placed its coordinates arbitrarily along the Twin Palm Road. We found fragmentary cranial and dental remains of one individual, and sampled a tooth (LP.01.01) which produced working aDNA (I10841) and is genetically male. A direct date places this individual in the 18th-20th centuries.

**Lusitu/Zimbe, Zambia**

*Southern Province, Latitude -16.13 Longitude 28.83*

*Primary reference: None*

This site is in the Lusitu area of Southern Province, and little information is available. Accession register information is as follows: *“Burial - Zimbe. 22m downstream of Kaiba* [should read Kariba]*, North fork?, in Baobab tree. Valley Tonga, C19, modern?*” An additional accession card reads "*Zimbe, LIA-modern, Burial Site. Fagan (Coll. 1961 BMF).*” Together these suggest the human remains were collected by Fagan, possibly in connection with his work at Ingombe Ilede, which is near Lusitu. The Ingombe Ilede report (Fagan et al. 1969:59, 168) [ref ^28^] mentions a 1961 survey in the appropriate area, as well as surface scatters of pottery at Lusitu, but makes no mention of a burial. We assigned estimated coordinates for the site. Remains are fragmentary and include the skull, pelvis, and long bones of a probable adult male. We sampled a tooth (LU.01.01), producing aDNA (I10832) from a genetic male, dated to the 18th-20th centuries.

**Mayburgh’s Farm, Zambia**

*Lusaka Province, Latitude -15.41 Longitude 28.28*

*Primary reference: None*

This site is in the Lusaka area, and little else is known about it; an alternative spelling may be Myburgh, a relatively common surname in Lusaka. The accession register notes that this is a “subsurface burial in cave graves,” assigned to “Bantu culture,” and notes the donor as R.P. Odendaal, who visited the Chipongwe site and collected human remains there (Clark & Toerien 1955) [ref ^4^]; however we have no basis to link Mayburgh’s Farm to Chipongwe. The human remains were wrapped in a 1951 newspaper with a note to Clark recording the location as “Mayburgh’s Farm, Nr. Lusaka, 12.3/24.17.” Here, we use Lusaka as approximate coordinates. We identified fragmentary remains of a maxilla and mandible, from which we sampled a tooth (MF.01.01), generating working aDNA (I10834) and a direct date falling in the first half of the 15th century.

**Moresby-White Farm, Zambia**

*Central Province, Latitude -14.46 Longitude 28.43*

*Primary reference: None*

The accession record notes that Moresby-White Farm is located at Broken Hill, Kabwe; we use this locality as coordinates. The accession register notes the burial is “Later Iron Age to modern.” Anderson’s (1961)[ref ^29^] report on Shimabala includes a short note written by Fagan that compares pottery from Shimabala with that of the Moresby-White site, in which he suggests that the latter dates to the 18th century. We found limb bones with large copper bangles, as well as a partial cranium with no teeth but both petrous portions intact; we note that more than one individual may be present in the fragmentary cranial remains. We sampled a petrous *in situ* using the cranial base drilling method (MW.01.01), producing working aDNA (I8387) from a genetic female, and a direct date falling in the late 17th-18th centuries, confirming Fagan’s instincts based on the material culture of this site and Shimabala.

**Mr. Bartlett’s Garden, Zambia**

*Lusaka Province, Latitude 15.41 Longitude 28.28*

*Primary reference: None*

There is no information available for this site, and we placed coordinates in Lusaka. The accession is associated with Chaplain, a midcentury official at the National Monuments Commission; the record reads “*Skeletal remains + 2 sherds, Mr Bartlett's Garden, Lusaka, ?modern*.” We found fragmentary remains of one adult probable male, with frequent red staining on the bones. We sampled the petrous bone (BG.01.01) of this individual, producing working aDNA (I10742) and confirming genetic male sex, but with insufficient collagen for a direct date

**Mtemankhokwe I, Malawi**

*Mangochi District, Southern Region, Latitude -14.5 Longitude 35.18*

*Primary reference: Juwayeyi 1991[ref* ^30^*]; Morris & Ribot 2006 [ref* ^31^*]*

This is a well-documented site in the Lower Shire Valley, not far from Nkudzi Bay. Juwayeyi (1991) excavated a Later IA cemetery here, and provided oral historical evidence suggesting a connection to present-day Nyanja-speaking Malawians. Juwayeyi (1991) noted strong similarities between Mtemankhokwe and Nkudzi Bay in terms of burial practices. His excavations recovered six individuals, all adorned with beads (including European trade beads), and in one case an ivory bangle, and accompanied by whole pots and iron tools. Morris and Ribot (2006) conducted bioarchaeological analysis of the six individuals (Mt1-6), curated at the Lilongwe Museum (now Museums and Monuments Department): five skulls (three attributed to males, two to females) and one mandible with no associated cranium (attributed to a female). All but one of these individuals (MTEM-1) could be located when visiting the collection for this study. We sampled the other five individuals for aDNA, via petrous bone (MTEM-2, -5, -6) or tooth (MTEM-3, -4). Both teeth and one petrous produced working aDNA (I6160, I6389, I6375), while another petrous produced low coverage results and was excluded (I6161), and another failed. Direct dates on each of the teeth fall between the late 17th and 20th centuries, confirming Juwayeyi’s (1991) indications of the site’s chronology, and its placement along trade networks that would have connected Mtemankhokwe I to the Swahili coast and Portuguese trading posts.

**Nansanzu School, Zambia**

*Southern Province, Latitude -17.85 Longitude 25.85*

*Primary reference: Vogel 1971[ref* ^27^*]*

This well-documented site is located in the Linda area of Livingstone and was excavated by Vogel (19710. In 1967, burials were discovered, along with pottery and grinding stones, during drain digging at the Nansanzu School. Vogel concluded that there were at least two distinct components, a village midden and a cemetery, and suggested that the burials might be intrusive and dating to the 19th century. Eight burials were uncovered, with five others said to have been removed during construction. These fragmentary burials, their positions, variable degrees of preservation, and associated grave goods are described in detail.

We found the fragmentary remains of at least three individuals commingled with faunal remains: two adult females, and one adult male. Labeling suggests these represent Burial V and possibly Burial II, but it is difficult to assign most remains to a burial, and it seems that many of the burials are missing. We sampled a tooth from remains clearly labeled Burial V (NZ.01.01), another unnumbered individual via a tooth (NZ.02.01), and a third unnumbered individual via an *in situ* petrous bone using cranial base drilling (NZ.03.01). The latter two individuals produced working aDNA (I10831, I8385) and were confirmed as genetic male and female, respectively. Direct dates on these two individuals are closely aligned and fall in the late 16th to early 17th centuries, substantially earlier than suggested by Vogel (1971).

**Ndonde, Zambia**

*Southern Province, Latitude -16.73 Longitude 27.37*

*Primary reference: Fagan 1978[ref* ^18^*]*

This site was accidentally discovered at Ndonde School, southeast of the IA site of Gundu, when human remains were found eroding out of an anthill. This led to investigation by Inskeep, who suggested that the remains were likely modern and perhaps related to a smallpox epidemic^18^. Fagan later conducted new excavations, documenting Early and Later/Recent IA cultural material, with charcoal dates in the middle first to early second millennium CE. He did not report additional burials, so the human remains cannot be easily associated with the material culture and stratigraphy documented by Fagan. We found five individuals previously named Ndonde A-E. We sampled a child (Burial B) via tooth (NDB.01), the near-complete skull of an adult via tooth (NDC.01) and three fragmentary crania of adults (Burials A, D, and E) via petrous bones (NDA.01, NDD.01, NDE.01). The four adult individuals produced working aDNA: Burial A (I10743), C (I10885), D (I10744), and E (I10745). Direct dates for these four burials are widely dispersed across the 10th-15th centuries, suggesting repeated use of this burial ground. The burials are at least broadly aligned with early second millennium dates for the site offered by Fagan (1978) based on Kalomo and Kangila pottery.

**Nkudzi (Nkhudzi) Bay, Malawi**

*Mangochi District, Southern Region, Malawi, Latitude -14.18 Longitude 35.01*

*Primary reference: Inskeep 1965[ref* ^32^*]; Morris & Ribot 2006 [ref* ^31^*]*

Nkudzi (also spelled Nkhudzi) Bay is a cemetery located along the western shores of what was then called Monkey Bay, at the southern end of Lake Malawi. Juwayeyi’s (2020)^33^ synthesis of regional oral traditions and archaeology provides context for understanding the cemetery, and he argues that it was used by the ancestors of Nyanja-speaking Malawians. Although this site is in Malawi, all but two of the individuals are curated at the LM. This site has been relatively well documented, first by Inskeep’s detailed excavation report (1965), later through Juwayeyi’s (1991)^30^ comparative analysis of the nearby cemetery of Mtemankhokwe I, and finally through bioarchaeological studies of the two individuals (NK1 and NK2) housed in Malawi, by Morris & Ribot (2006), and two individuals housed in Livingstone (accession 6709) by Bräuer and Rösing (1989)^34^. Most of the Nkudzi Bay skeletal collection in Livingstone remains unstudied.

The cemetery was discovered in 1957 when burials were being eroded by the lake. Inskeep (1965) noted that part of the site was already underwater, and human remains and grave goods could be found ~20m off the shoreline; two skulls were collected from the lake bed (accession 6709). Over eight days in 1958, Inskeep exposed 9.5 m^2^ in an area of a sandy peninsula where the cemetery was still mostly intact, documenting (but not fully excavating) 12 burials (ref ^32^). This rapid excavation was made more challenging by superimposition of burials, lack of clear soil color changes or pits, and problems of erosion and scattering of human remains. Inskeep acknowledged that the minimum number of individuals might exceed 12, and that it was challenging to establish a relative chronology, or to associate grave goods with specific burials, except for Burials I, VI, X, and XI. Material culture included abundant glass beads, imported ceramics, and other goods suggesting the cemetery was in use during the 18^th^-19th century (ref ^32^).

The two individuals curated in Blantyre cannot be clearly linked to the numbered Burials I-XII in Inskeep (ref ^32^). At the LM, accession number 6709/SK1 corresponds to the two crania from the lake bed. Accession number 7016 (SK30-SK33) is the larger group of remains from the 1958 excavations. The register notes 10 human skeletons of various ages. As of 2017, remains at LM are distributed across three boxes, in bags usually containing more than one individual and labelled with context information. Almost no skeletal remains are directly labelled, and labels do not link back to Inskeep’s (1965) burial numbers. The remains are fragmentary, incomplete and commingled, but generally well preserved, particularly teeth. Many elements were treated with consolidant. There was also prior sampling via femur cross-sectioning.

We struggled to link these remains to the burials described by Inskeep (1965). Burials IV and VI were not collected, and Burial XI was not fully excavated and are unlikely candidates. Burials II and VIII-X, all adults, may be present at the LM among the commingled remains, but are only represented by postcrania, making it challenging to separate unique individuals. This leaves Burials I, III, V, VII, and XII as most likely to have cranial remains in the LM. However, we documented at least 16 individuals represented by cranial remains: eight non-adults ranging from newborn to teen, seven adults, and one individual with a small mandible and incomplete dentition that could either be an older child or adult. Six individuals can be linked back to Burials I, II, II, V, VII, and XII, using age, dental development, and/or skeletal elements present.

In this study, we sampled all 16 individuals identified via morphology at the LM, two of which were genetically the same individual, despite different contexts. Additionally one of the two crania at the Malawi National Repository in Blantyre was sampled via the left petrous (labeled MAD45, which we infer based on sex is NK2). Direct dates on eleven individuals fall within the 17th-20th centuries; two individuals could date to as early as the 15th and 16th centuries. Due to wiggles in postindustrial calibration curves, there is low dating resolution for most individuals from this site. Broadly, many dates are consistent with Inskeep’s estimate of the 18th-19th century.

**Olympia Park, Zambia**

*Lusaka Province, Latitude -15.40 Longitude 28.30*

*Primary reference: Phillipson 1970 [ref* ^35^*]*

This site was discovered during construction of the National Assembly building at Olympia Park Hill in Lusaka. Phillipson (1970) reported that several graves were found in foundation trenches, and that they were associated with pottery similar to that found in topsoil at the nearby site of Twickenham Road, and therefore thought to be recent. No further information is available, but the burials were found relatively intact and heavily consolidated in the LM. Two of these were labeled as Olympia Park D and E; presumably the other individuals we found represent Burials A, B, and C. However we identified at least six individuals, all young adults except for the infant (Olympia Park E). We collected samples from all six: an ulna fragment from a young adult probable male (OP.01.01); a tooth from a young adult (OP.02.01); the isolated petrous of the Olympia Park E infant (OP.03.01); a tooth from another young adult (OP.04.01); a mandibular tooth (OP.05.01) and the *in situ* petrous of a cranium associated with Olympia Park D (OP.06.01). The mandible and cranium likely represent two distinct individuals. We obtained working aDNA from the Olympia Park E infant (OP.03.01, I10746), one of the young adults (OP.04.01, I8990), and the Olympia Park D adult petrous (OP.06.01, I8389). The latter two individuals produced direct dates falling in the late 17th-20th and late 17th-18th centuries, respectively, suggesting one or both are earlier than Phillipson (1970) suspected.

**Shimabala, Zambia**

*Lusaka Province, Latitude -15.65 Longitude 28.23*

*Primary reference: Anderson 1961[ref* ^29^*]*

A series of burials was discovered in 1959 during blasting at the Shimbala Quarry south of Lusaka, about a mile from the Chipongwe Caves. In a single vertical fissure, F.V. Anderson (1961) documented the remains of at least 35 and perhaps as many as 50 individuals, almost exclusively adults, deposited continuously over more than 5 meters of depth. Despite her pleas to the quarry managers to stop work, she was initially ignored, and many human remains were destroyed and context lost. Even once Anderson was allowed to conduct excavations, it remained exceedingly challenging to work from a high and narrow ledge, let alone to adequately document the remains.

An initial bioarchaeological assessment was made by J. Hiernaux (in Anderson 1961), who noted that the remains, representing at least 33 individuals, were so fragmentary that no skull could be reconstructed, nor could sexes be determined. Hiernaux noted some remains were slightly burnt. Anderson interpreted this as a “mass burial,” in which people in their prime age were deposited without ceremony, nor with weapons, and the charred skeletal material might indicate, in her view, a ritualistic and/or cannibalistic sacrifice, or alternatively, an epidemic. Anderson (1961) consulted with J.D. Clark and B.M. Fagan on material culture, which included shell beads and copper and iron bangles, potsherds from at least 20 vessels, and just four glass beads interpreted as evidence for limited contact with European trade networks. Fagan noted similarities between the pottery of Shimabala and Moresby-White Farm. Both Clark and Fagan independently concluded that a late 17th to early 18th century date might be appropriate for the site.

At the LM, we found the remains of at least nine individuals; the material was so fragmentary and commingled (including with faunal remains) that it was difficult to identify additional individuals; labels indicated that many of these remains belonged to a “Shi V Burial,” but this burial number clearly includes many individuals. To avoid double-sampling, we selected the right petrous portion of five individuals, and selected non-repeating teeth from another four individuals. Seven of these nine individuals produced working aDNA (I8871, I8872, I8985, I10716, I10717, I10718, I8993), five genetic males and two genetic females. Only four individuals had sufficient collagen for radiocarbon dating. Calibrated dates from three of these individuals cluster in the late 11th to early 13th centuries, much earlier than the tentative chronology offered by Clark and Fagan. A fourth individual dates to the 17th to 20th centuries.

**Sikalongo, Zambia**

*Southern Province, Latitude -16.97 Longitude 27.23*

*Primary reference: Musonda 1987[ref* ^36^*]*

This site is located in Choma District, and its coordinates are approximate. Human remains were accidentally discovered in a trash pit in 1986, leading to excavations by Musonda (1987). He documented a single burial at 1.5 m below surface, which he attributed to a relatively tall female, accompanied by pottery and a pipe. Nearby at Simulombwa Hill, he uncovered the remains of a village with six wattle-and-daub huts. Musonda (1987:42) suggested the burial might date to the Later IA, between the 11th and 18th centuries, based on associated pottery. He further drew upon ethnographic information on present-day burial traditions, and Tonga oral traditions, to provide additional interpretation. Notably, Musonda was informed that in previous digging for house and water pipe construction at Sikalongo, additional burials had been found with copper crosses and abundant pottery, recalling the burials of Ingombe Ilede, c. 160 km away. No further information is available about these burials or the whereabouts of the associated material culture.

In the LM collections, we found cranial vault fragments and postcrania consistent with a single individual, possibly adolescent or young adult, and possibly male. We sampled this individual via the petrous bone (SL.01.01), which produced working aDNA (I10741), confirmed male genetic sex, and gave a direct date in the 18th-20th centuries.

**Silverest (Silver Rest) Farm, Zambia**

*Lusaka Province, Latitude -15.37 Longitude 28.50*

*Primary reference: none*

The accession register for this site notes that it is east of Lusaka, is “*Iron Age? Recent*”, and mentions a healed wound on the left temple of the burial, which was given to the museum by Mr. P. Steyer. The site, written as “Silver Rest” in the register, is likely located at the present-day “Silverest Farm” east of Lusaka, and we have used these coordinates as an approximate location. Although a file (File 730/II/86) is referred to in the accession register, we were unable to locate it, and therefore we lack further information on this site. A single individual was represented by its cranium; we sampled this individual via the petrous bone (SV.01.01), which produced working aDNA (I10739) from a genetic male, directly dated to the 19th to early 20th centuries.

**Simbusenga, Zambia**

*Southern Province, Latitude -17.58 Longitude 25.58*

*Primary reference: Vogel 1975[ref* ^37^*]; for burials, Murphy 1996, 1999[ref* ^16,38^*]*

This is a well-documented site in Kalomo District, c. 35 northwest of Victoria Falls, excavated by Vogel in 1967 and 1969, and reported in detail in Vogel (1975). Vogel documented the remains of repeated village occupations with hiatuses, and a series of 18 charcoal dates place the site’s chronology in the Later IA, between c. 750-1575 CE. He noted similarities in material culture between Simbusenga and the sites of Sinde Mission and Nansanzu School. Material culture included Kalomo pottery, iron implements, clay figurines, pipes, and lip plugs.

Vogel excavated nine burials, accompanied by copper bracelets in some cases, as well as shell, ivory, and glass beads, and bark cloth. The chronology of the burials is based on associated radiocarbon dates and stratigraphic position. Burial 1 was thought to be intrusive from the Linda Phase (recent), while Burials 2-9 were attributed to c. 1100-1500 CE (per Vogel 1975)^37^ or c. 1400-1500 CE (per Murphy 1996)^16^. Unusually for Zambian IA sites, these burials have been well documented by a bioarchaeologist, including paleopathological and stable isotope analyses^16,38^. Murphy analyzed Burials 1-8, noting that Burial 9 was young and largely destroyed by ants^37^ and was not identifiable at the LM. All except Burial 1 were fragmentary. She recorded polydactyly in Burial 5, and pathological conditions in other burials.

We were able to identify all eight burials despite their fragmentary state, thanks to prior organization by either Murphy or S.K. McIntosh, who inventoried the site in 2017. We collected a sample from each individual, two of which produced working aDNA (Burial 7, I10079; Burial 8, I8378), from genetic females. This low success rate may be informative about the preservation conditions or age of the skeletons; the apparently modern Burial 1 did not produce aDNA. The two successful samples were directly dated to the late 13th through 14th centuries.

**Sinachisingili (Gwembe Valley), Zambia**

*Southern Province, Latitude -17.43 Longitude 27.47*

*Primary reference: none*

This site is located in the Gwembe Valley and is one of the sites that was documented during excavations ahead of the Kariba Dam project in the Middle Zambezi River; the site is now flooded. Little information is available about the burial: both Clark and Inskeep are listed on the accession card, which reads “*Grave. Iron Age. Exc. A. 21-8-57. See field notes*.” No field notes could be found. LSA and Early IA artifacts were documented at this locality, according to the NHCC catalog. We found a fragmentary, partial cranium and mandible. We sampled an isolated petrous bone (SG.01.01), which produced working aDNA (I8869) from a genetic female. There was insufficient collagen in this sample for a radiocarbon date.

**Sinde Mission Orange Grove, Zambia**

*Southern Province, Latitude -17.67 Longitude 25.83*

*Primary reference: Inskeep 1962[ref* ^39^*]*

This site is located ~23 km north of Livingstone; it is distinct from the Sinde site recorded by Vogel (1971)[ref ^27^]. Inskeep (1962) noted that pottery collected around Sinde Mission resembled the earliest pottery from Kalundu Mound. In 1957, he was alerted to human remains discovered by digging irrigation pits, and visited the site with J.D. Clark. They noted damaged human remains representing multiple burials, two complete pots, carinated bowls, iron implements, and a worked bird bone tube. A local informant suggested the cemetery was recent, so they decided not to excavate further. Missionaries later reported to Inskeep finds that turned up in agricultural work, including a copper cross (on the surface), pipe bowls, cowrie shells, and 1,758 glass beads. Based on his informant’s testimony, Inskeep (1962) concluded the cemetery dated to the turn of the 20th century, despite the earlier material culture.

At the LM we found the commingled and fragmentary remains of three individuals. Two are associated with Clark and Inskeep’s accession (5723), one with a later accession by a Mrs. Boddell (6041). We sampled two individuals via their petrous bones (5723/SK19, SM.01.01; and 6041/SK6, SM.02.01), and a third via an incisor (5723/I27, SM.03.01). The petrous bones produced working aDNA (I10727 and I10728) from two genetic males, only one of which had sufficient collagen for radiocarbon dating. This date falls in the late 17th to 18th centuries, earlier than suggested by Inskeep’s informant, and consistent with the material culture.

**Twin Rivers Kopje, Zambia**

*Lusaka Province, Latitude -15.56 Longitude 28.13*

*Primary reference: Clark & Brown 2001[ref* ^40^*].*

This is a well-documented site southwest of Lusaka. It is a kopje (inselberg or hill) with shelters, sinkholes, and a shallow cave, best known for its Late Pleistocene deposits^40,41^. However it also bears more recent occupations, including a single Recent IA burial in the shallow cave, in which Clark & Brown (2001:308) documented an adult and child associated with “shallow bowls, metal ornaments and shell and glass beads.” No further information was provided. We found fragmentary and commingled remains of at least four and likely five individuals. We identified either one or two adults (TR.01.01 and TR.05.01), from which we sampled a petrous bone and a canine, respectively; and at least three subadults which were each sampled via an incisor: a perinatal infant (TR.03.01), at least one young child (TR.02.01), and an adolescent (TR.04.01). Three of these samples produced working aDNA: a genetically male young child (TR.02.01/I8981), dating to the late 12th-13th century; and a genetically female adolescent (TR.04.01/I8983) and genetically female adult (TR.01.01/I8870) which both date to the first half of the 15th century, suggesting this reportedly single grave represents multiple burial events.

**Water Tower, Livingstone, Zambia**

*Southern Province, Latitude -17.85 Longitude 25.87*

*Primary reference: none*

This site is located in Livingstone, and the catalog card notes it was found ½ mile north of the water tower, on the south side of the Great North Road, and was deposited in July 1959. No further information could be found. We found the remains of a single adult, probably male. We sampled the petrous via cranial base drilling (WT.01.01), and this produced working aDNA (I8388) from a genetic male, which directly dated to the late 17th to 20th centuries.

**Zambezi River Bank, Zambia**

*Southern Province, Latitude unknown Longitude unknown*

*Primary reference: none*

This site is sparsely documented, and we cannot assign even approximate coordinates. The catalog for this accession number (7987, SK 43), reads “*Zambezi River Bank. RIA/modern. Burial site*,” but no catalog card could be found, and we could not find further records of this site in the museum or in literature review. While Vogel (1973)^26^ does record a 19th century burial at the Zambezi Farm site, we have no reason to suspect these are the same. We found the remains of at least two individuals: a juvenile represented by fragmentary long bones, and a young adult male represented by maxilla and mandible. Another bag labeled “ZB floating bones” contained a petrous portion and additional cranial fragments. We sampled an incisor and a petrous from what we thought might be two distinct adults (ZB.01.01/ZB.02.01), but these are the same individual (I10740), a genetic male, directly dating to the 18th-20th centuries.

**Radiocarbon Dating and Revised Chronologies**

All samples from the LM collections that produced readable aDNA, plus two samples from Mtemankhokwe in Malawi, were submitted to the Pennsylvania State University (PSU) Radiocarbon Laboratory for radiocarbon dating via accelerator mass spectrometry (AMS) (**Supplementary Table 2**). For the collagen samples (all those except from Mtemankhokwe), carbon and nitrogen concentrations and stable isotope ratios were measured at the Yale Analytical and Stable Isotope Center. We calibrated all dates using OxCal v.4.4^42^ with the SHCal20 curve^43^, and we provide the 2σ range and median values in **Supplementary Table 2**. Plots of the calibrated dates are found below for sites in Zambia and Malawi. Since a majority of dates in this paper fall within the last three centuries, they are subject to the ‘Suess effect’ in which the burning of fossil fuels since the 18th century has diluted atmospheric ^14^C. While the calibration curve takes this into account, wiggles in the curve lead to low resolution for 18th-20th century samples.

Despite these uncertainties with recent dates, a few observations can be made. First, this represents a substantial effort at direct AMS dating human remains from sites that either had no available chronology, or a series of indirect dates often predating widespread use of AMS. These new dates benefit repositories that do not have the resources for widespread dating efforts, and whose curators wished to know more of the collections under their care. Second, the revised chronologies for many sites underscore the importance of reassessing regional chronologies via legacy collections, and will hopefully generate new research questions for future scholars to do so. Recent work in Zambia, for example, has shown that several sites have more recent deposits than was previously appreciated^13,44,45^.

The vast majority of individuals in our study fall within the 17th to 20th centuries, as was expected given scant contextual information and indications, in many cases, that these were recent burials. However at several sites, burials are earlier than was expected either based on local information, material culture or on the apparently intrusive position of graves. At both Kanyankunde Rockshelter and Mayburgh’s Farm, previously undated burials were directly dated to the first half of the 15^th^ century, providing these sites’ only chronological anchors. At Chipongwe Cave, where burials initially thought to date to the late 19^th^ century may be up to two centuries earlier. Similarly, at Nansanzu School, burials thought to be intrusive and recent instead date to the late 16th to early 17th centuries, and at Ndonde, burials thought to be from a modern smallpox epidemic instead fall within the 10th-15th centuries; for both sites, this suggests closer than expected ties between the burials and IA material culture found there. At Olympia Park and at Sinde Mission, burials thought to be recent also date to the 17^th^-18^th^ centuries.

At several other sites, direct dating revealed complex site formation histories and repeated use of burial grounds. This is the case at Kalala Island, where direct dates on six individuals suggest repeated use of the island as a burial ground at various points falling between the 14^th^ and 20^th^ centuries. While Derricourt (1985)^17^ was correct that some burials were intrusive and recent, there are others that may be more clearly relatable to the IA material culture at the site. Similarly, at Nkudzi Bay, most direct dates overall confirm Inskeep’s (1965)[ref ^32^] interpretation of the site as falling within the 18^th^-19^th^ centuries, but at least two individuals were buried as early as the 15^th^ and 16^th^ centuries. At Shimabala, direct dates on four individuals show that there is no single mass burial event, as originally suggested by Anderson (1961)^29^, and that at least three out of the dozens of individuals buried at the site date to the late 11th to early 13th centuries, far earlier than was suspected. At Twin Rivers Kopje, an undated Recent IA burial had far more individuals in it than previously reported, and represented multiple burial events falling in the late 12th-13th centuries and early 15^th^ century.

Several sites in our dataset would benefit from additional chronological reassessment. No new dates were generated for the Ingombe Ilede burials, and this is an area where additional attention to direct dating would be especially valuable. It would also make sense to further date sites with multiple inhumations, as at Chipongwe, Shimabala, Simbusenga, and Twin Rivers Kopje, including dating individuals that did not produce aDNA. Finally, it would be informative to have additional dates from Malawian sites in our dataset, especially Chipakusa and Dzimbiri.

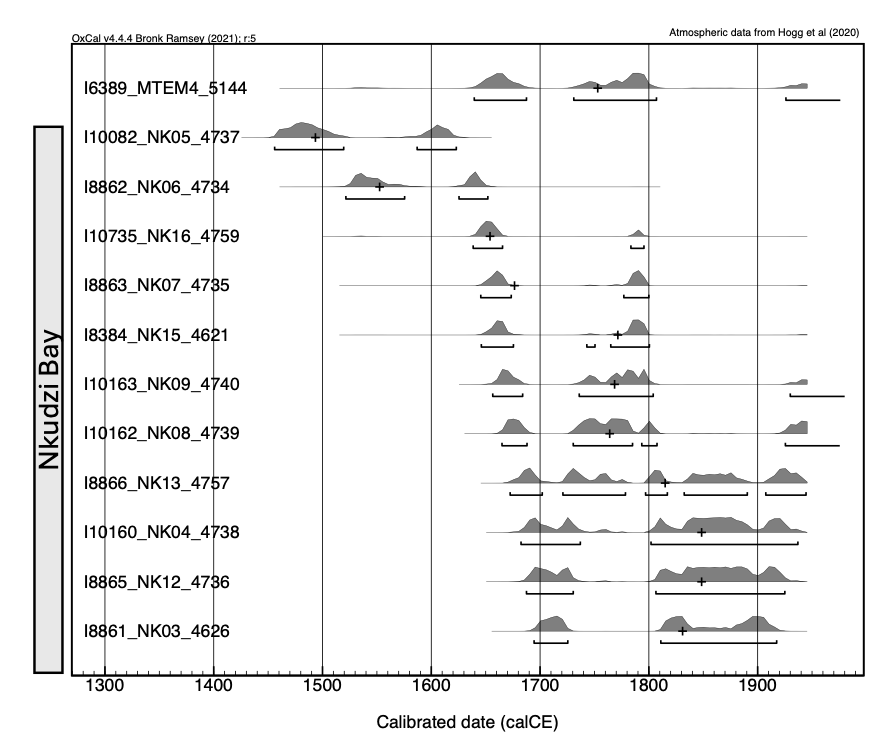

**Figure S2.** Calibrated radiocarbon dates from sites in Malawi: Mtemankhokwe (top) and Nkudzi Bay (all others), ordered chronologically within each site. Sample codes combine Individual ID, Sample ID, and PSUAMS lab number. Note the lack of resolution in dates spanning the late 17th-20th centuries.

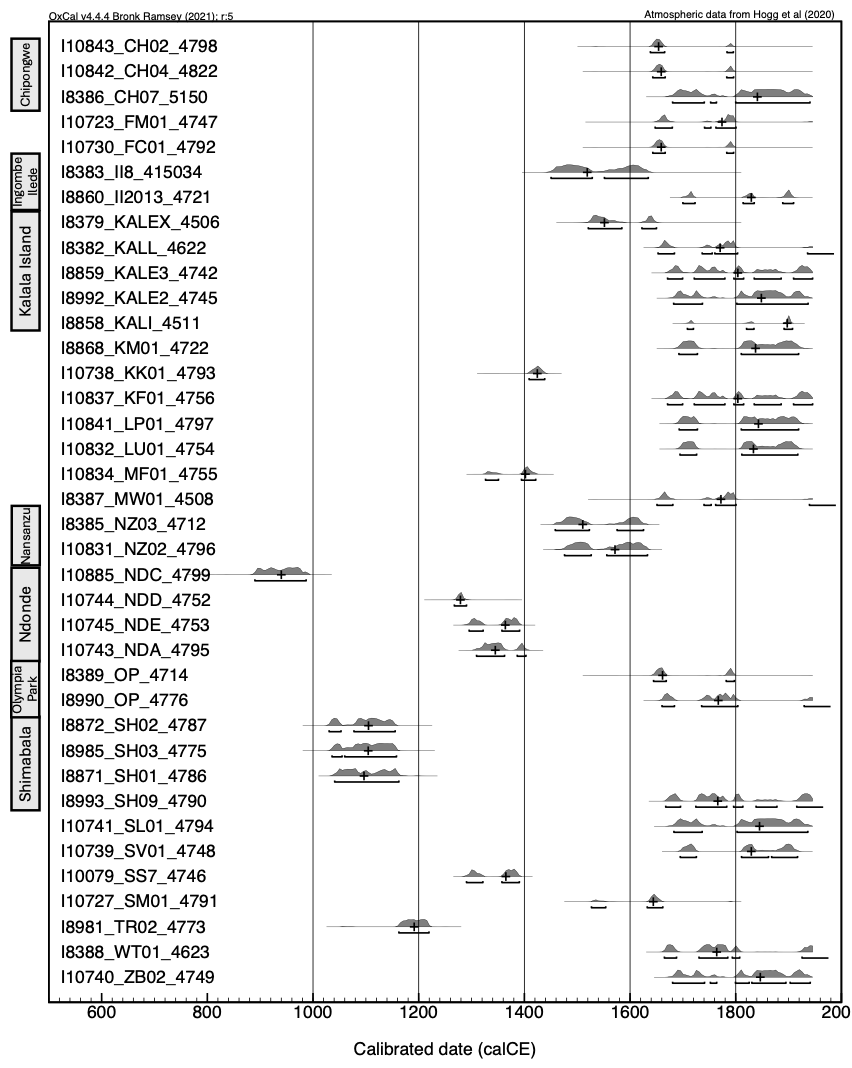

**Figure S3.** Calibrated radiocarbon dates from sites in Zambia, ordered alphabetically by site and chronologically within each site. Sites with several radiocarbon dates are called out in gray bars to the left. Sample codes combine Individual ID, Sample ID, and PSUAMS lab number. Note the lack of resolution in dates spanning the late 17th-20th centuries.

1.	Skoglund, P. *et al.* Reconstructing Prehistoric African Population Structure. *Cell* **171**, 59-71.e21 (2017).

2.	Lipson, M. *et al.* Ancient DNA and deep population structure in sub-Saharan African foragers. *Nature* **603**, 290–296 (2022).

3.	Dussubieux, L., Welling, M., Kaliba, P. & Thompson, J. C. European Trade in Malawi: The Glass Bead Evidence. *Afr. Archaeol. Rev.* **40**, 377–396 (2023).

4.	Clark, J. D. & Toerien, M. J. Human skeletal and cultural material from a deep cave at Chipongwe, Northern Rhodesia. *South Afr. Archaeol. Bull.* **10**, 107–116 (1955).

5.	Dart, R. A. & del Grande, N. The ancient iron smelting cavern at Mumbwa. *Trans. R. Soc. South Afr.* **19**, 379–427 (1931).

6.	Miller, S. E. F. *The Nachikufan Industries of the Later Stone Age in Zambia*. (1969).

7.	Clark, J. D. *The Stone Age Cultures of Northern Rhodesia: With Particular Reference to the Cultural and Climatic Succession in the Upper Zambezi Valley and Its Tributaries*. 157 (The South African archaeological society, Claremont, Cape, 1950).

8.	Fortes-Lima, C. A. *et al.* The genetic legacy of the expansion of Bantu-speaking peoples in Africa. *Nature* **625**, 540–547 (2024).

9.	Villiers, H. & Fatti, L. P. Antiquity of the Negro. in *Para conocer al hombre: homenaje a Santiago Genovés a 33 años como investigador en la UNAM* 299–312 (Universidad Autónoma de México, Instituto de Investigaciones Antropológicas, México, 1990).

10.	Kaiser, T. M., Seiffert, C. & Truluck, T. The speleological potential of limestone karst in Zambia (Central Africa) – a reconnaissance survey. *Cave Karst Sci.* **25**, 23–28 (1998).

11.	Chaplin, J. H. A preliminary account of Iron Age burials with gold in the Gwembe Valley, Northern Rhodesia. in *Proceedings of the 1st Federal Science Congress, 1960* 397–406 (1962).

12.	Fagan, B. M., Phillipson, D. W. & Daniels, S. G. H. *Iron Age Cultures in Zambia (Dambwa, Ingombe Ilede and the Tonga)*. vol. 2 (London, 1969).

13.	McIntosh, S. K. & Fagan, B. M. Re-dating the Ingombe Ilede burials. *Antiquity* **91**, 1069–1077 (2017).

14.	Gibbon, V. E., Gallagher, A. & Huffman, T. N. Bioarchaeological Analysis of Iron Age Human Skeletons from Zambia. *Int. J. Osteoarchaeol.* **24**, 100–110 (2014).

15.	Phillipson, D. W. & Fagan, B. M. The date of the Ing’ombe Ilede burials. *J. Afr. Hist.* **10**, 199–204 (1969).

16.	Murphy, K. A. *The Skeletal Elements of the Iron Age in Central and Southern Africa: A Bioarchaeological Approach to the Reconstruction of Prehistoric Subsistence*. (1996).

17.	Derricourt, R. M. *Man on the Kafue: The Archaeology and History of the Itezhitezhi Area of Zambia*. (Ethnographica, 1985).

18.	Fagan, B. M. Gundu and Ndonde, Basanga and Mwanamaimpa. *Azania J. Br. Inst. East. Afr.* **13**, 127–134 (1978).

19.	Fagan, B. M. *Iron Age Cultures in Zambia (Kalomo and Kangila)*. vol. 1 (Chatto & Windus National Museum of Zambia, 1967).

20.	Gear, H. S. A Boskopoid skeleton from Kalomo, Northern Rhodesia. *Bantu Stud.* **2**, 217–231 (1923).

21.	Schepers, G. W. H. A fossilised human mandible from Kopje Alleen, western Transvaal. *South Afr. J. Sci.* **32**, (1935).

22.	Clark, J. D. Further excavations (1939) at the Mumbwa caves, Northern Rhodesia. *Trans. R. Soc. South Afr.* **29**, 133–201 (1942).

23.	MacCrae, F. B. & Lancaster, D. G. Stone Age sites in Northern Rhodesia. *Man* **37**, 62–64 (1937).

24.	Phillipson, D. W. *The Prehistory of Eastern Zambia*. (Nairobi, 1976).

25.	Fagan, B. M. & Phillipson, D. W. Sebanzi: The Iron Age sequence at Lochinvar, and the Tonga. *J. R. Anthropol. Inst. G. B. Irel.* **95**, 253–294 (1965).

26.	Vogel, J. Some Early Iron Age sites in southern and western Zambia. *Azania* **8**, 25–54 (1973).

27.	Vogel, J. *Kamangoza*. (1971).

28.	Fagan, B. M., Phillipson, D. W. & Daniels, S. *Iron Age Cultures in Zambia*. vol. 2 (Chatto and Windus, London, 1969).

29.	Anderson, F. V. The Shimabala mass burial. *South Afr. Archaeol. Bull.* **16**, 144–147 (1961).

30.	Juwayeyi, Y. M. Late Iron Age burial practices in the southern Lake Malawi area. *South Afr. Archaeol. Bull.* 25–33 (1991).

31.	Morris, A. G. & Ribot, I. Morphometric cranial identity of prehistoric Malawians in the light of sub-Saharan African diversity. *Am. J. Phys. Anthropol.* **130**, 10–25 (2006).

32.	Inskeep, R. R. *Preliminary Investigation of a Proto-Historic Cemetery at Nkudzi Bay, Malawi*. (1965).

33.	Zubieta, L. F. Yusuf M. Juwayeyi. 2020. Archaeology and oral tradition in Malawi: origins and early history of the Chewa. New York: Boydell & Brewer; 978-1-84-701253-1 hardback £60. *Antiquity* **95**, 1352–1354 (2021).

34.	Braüer, G. & Rösing, F. W. Human biological history in southern Africa. in *Rassengeschichte der Menschheit. 13. Lieferung. Afrika II: Südafrika. Schwidetzky I, editor.* 7–137 (R.Oldenbourg, Munich, 1989).

35.	Phillipson, D. W. Excavations at Twickenham Road, Lusaka. *Azania J. Br. Inst. East. Afr.* **5**, 77–118 (1970).

36.	Musonda, F. B. The significance of pottery in Zambian Later Stone Age contexts. *Afr. Archaeol. Rev.* **5**, 147–158 (1987).

37.	Vogel, J. *Simbusenga: The Archaeology of the Intermediate Period of the Southern Zambian Iron Age*. (Oxford University Press, 1975).

38.	Murphy, K. A. A prehistoric example of polydactyly from the Iron Age site of Simbusenga, Zambia. *Am. J. Phys. Anthropol.* **108**, 311–319 (1999).

39.	Inskeep, R. R. Some iron age sites in Northern Rhodesia. *South Afr. Archaeol. Bull.* **17**, 136–180 (1962).

40.	Clark, J. D. & Brown, K. S. The Twin Rivers Kopje, Zambia: Stratigraphy, Fauna, and Artefact Assemblages from the 1954 and 1956 Excavations. *J. Archaeol. Sci.* **28**, 305–330 (2001).

41.	Clark, J. D. Human Behavioral Differences in Southern Africa During the Later Pleistocene. *Am. Anthropol.* **73**, 1211–1236 (1971).

42.	Ramsey, C. B. Bayesian Analysis of Radiocarbon Dates. *Radiocarbon* **51**, 337–360 (2009).

43.	Hogg, A. G. *et al.* SHCal20 Southern Hemisphere Calibration, 0–55,000 Years cal BP. *Radiocarbon* **62**, 759–778 (2020).

44.	McKeeby, Z. Mapping the Iron Age in Southern Africa: Magnetometry at two Iron Age villages in Western Zambia. *J. Archaeol. Sci.* **163**, 105937 (2024).

45.	Goldstein, S. T. *et al.* Revisiting Kalundu Mound, Zambia: Implications for the Timing of Social and Subsistence Transitions in Iron Age Southern Africa. *Afr. Archaeol. Rev.* **38**, 625–655 (2021).
