## Supplementary figures and images for "The south Congo Basin was critical to Bantu settlement of south central Africa"

### Supplementary Figures 1

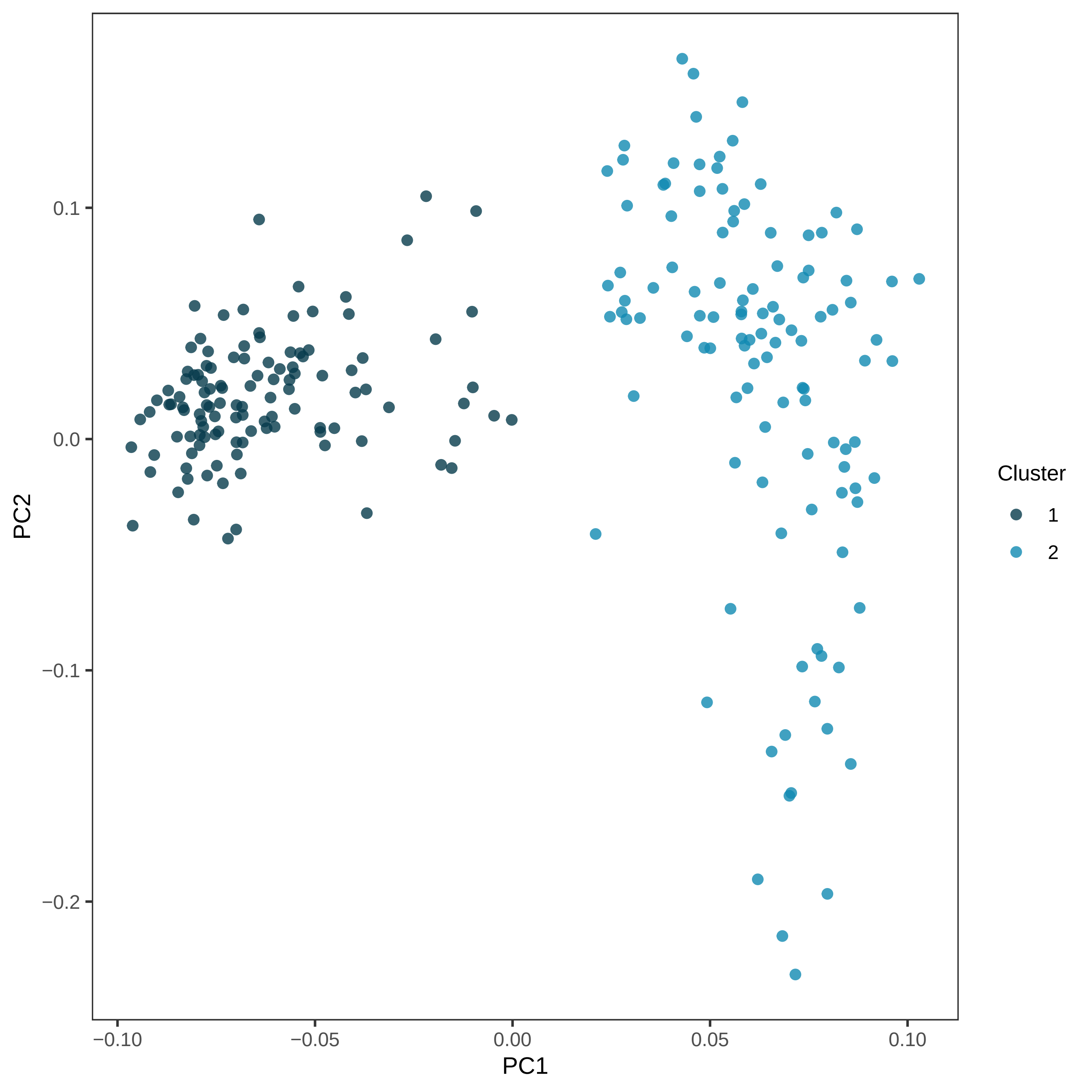

### Supplementary Figures 2

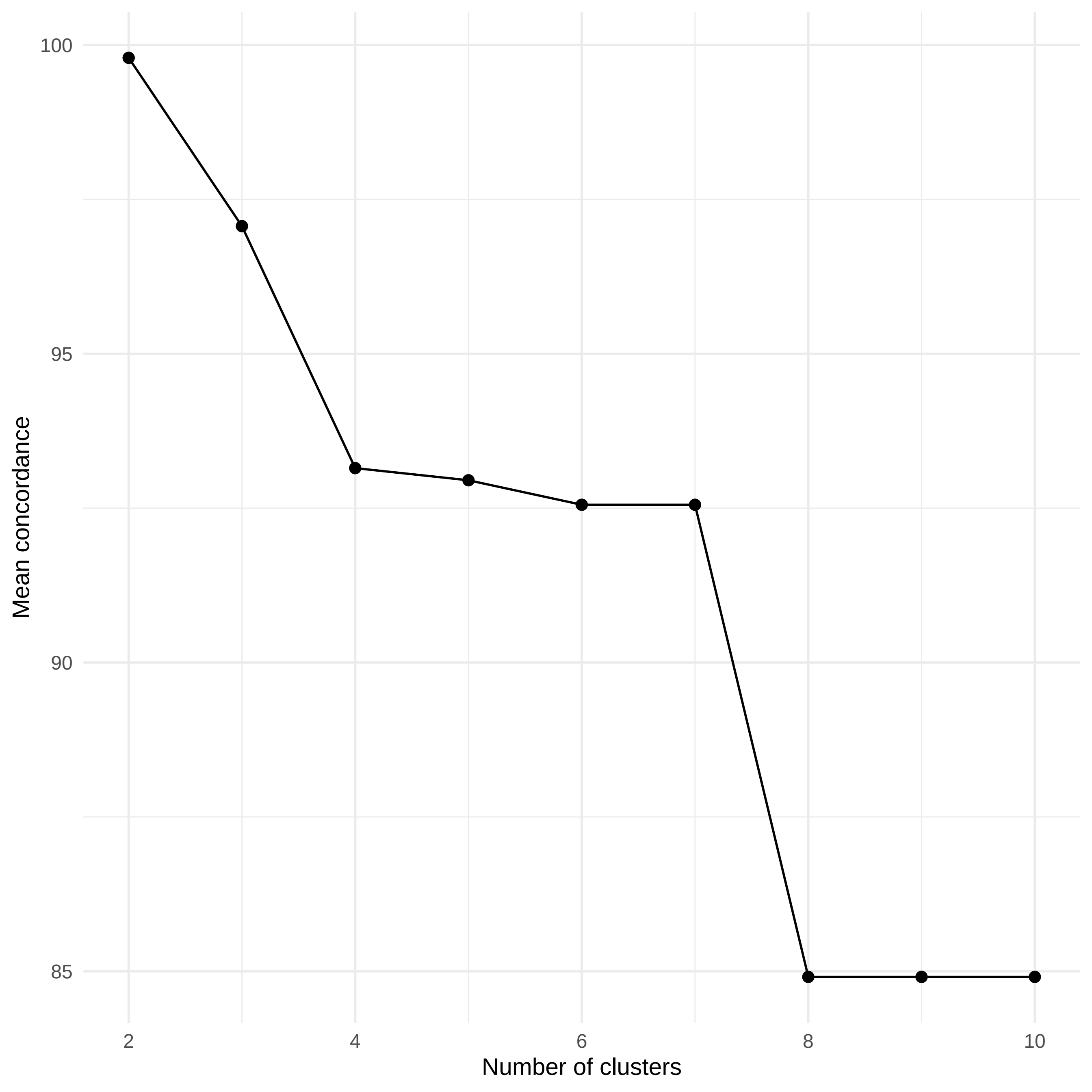

### Supplementary Figures 3

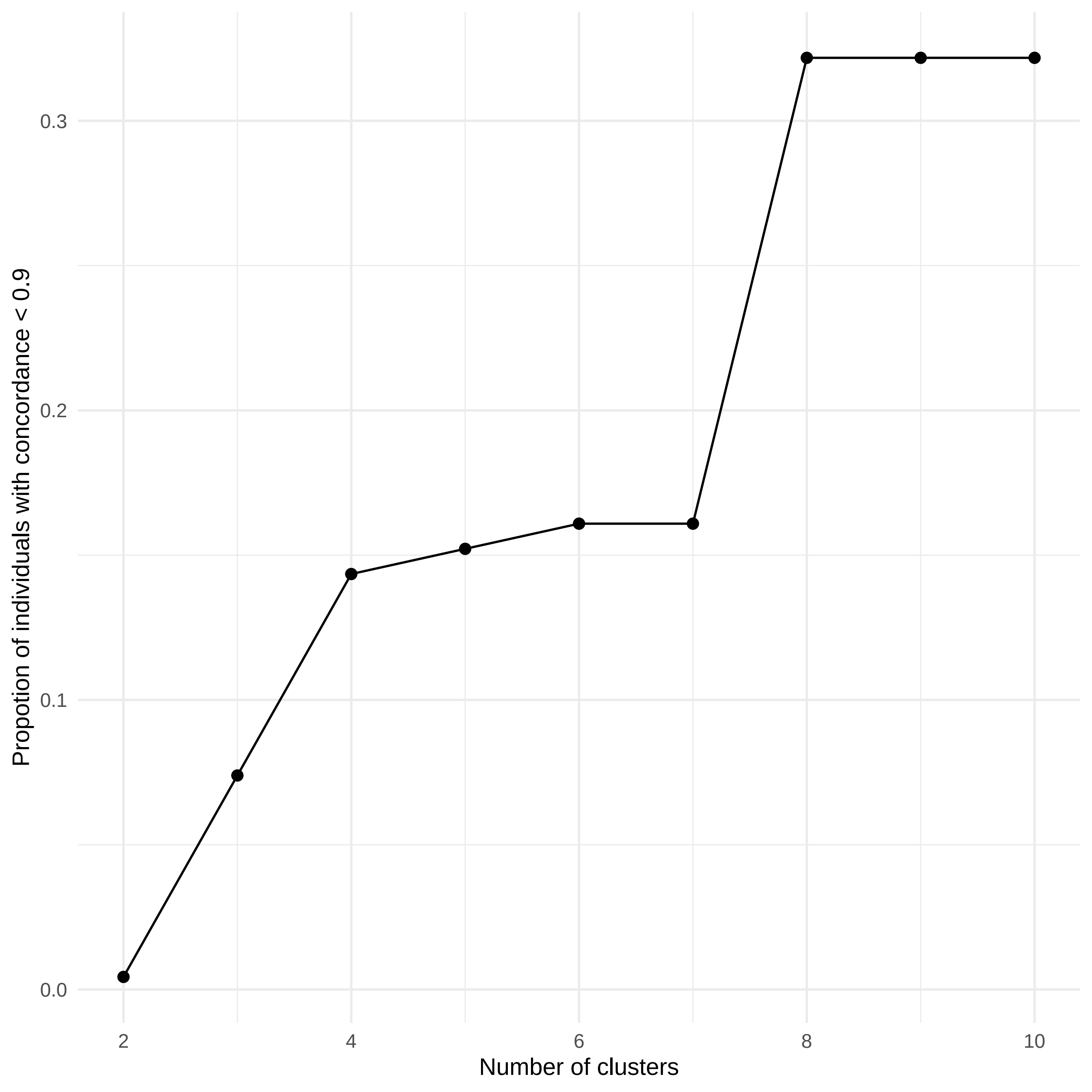

### Supplementary Figures 4

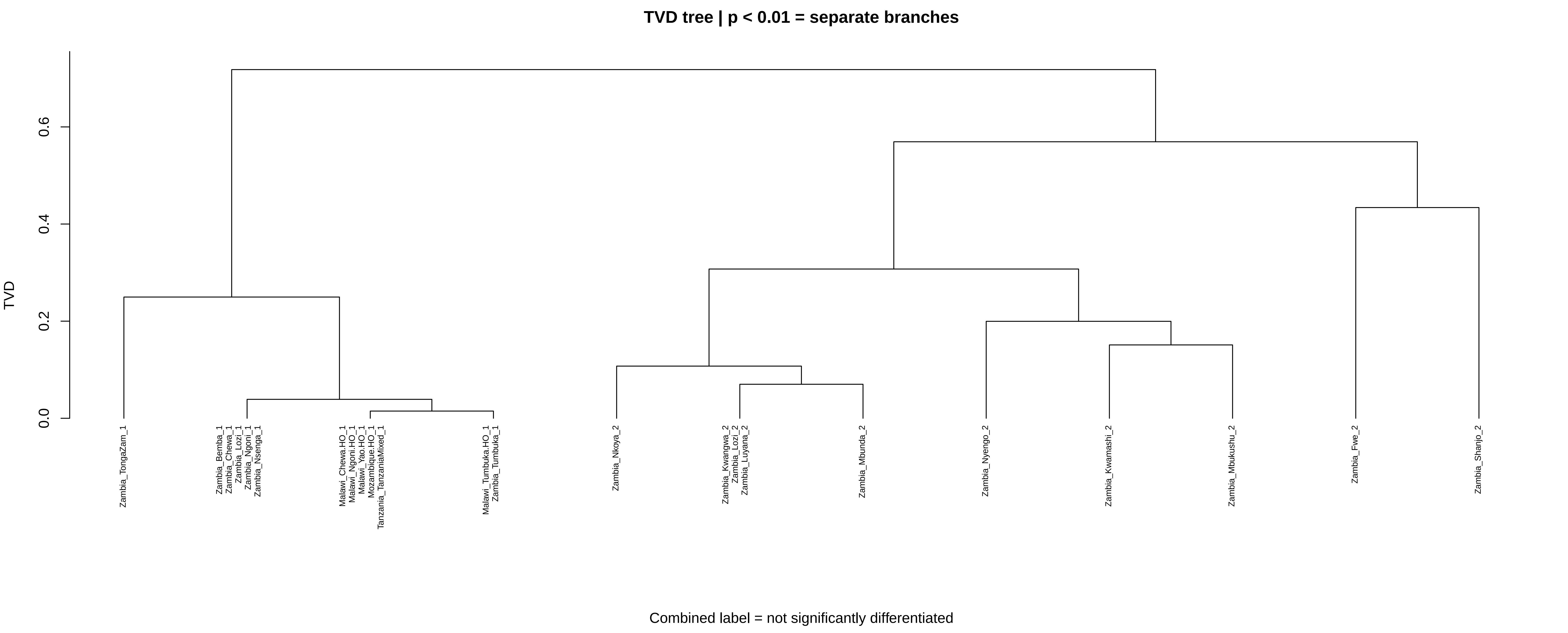

### Supplementary Figures 5

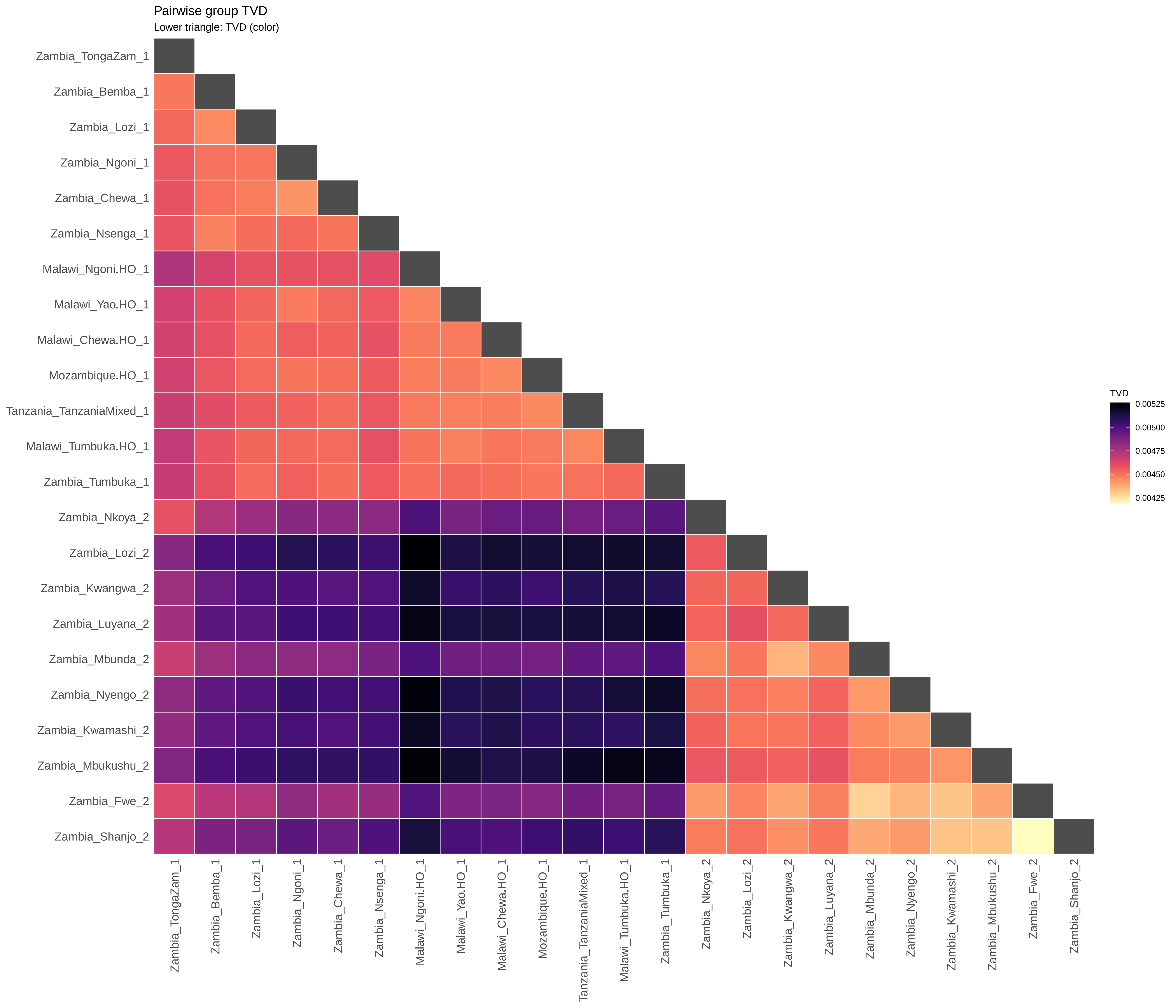

### Supplementary Figures 6

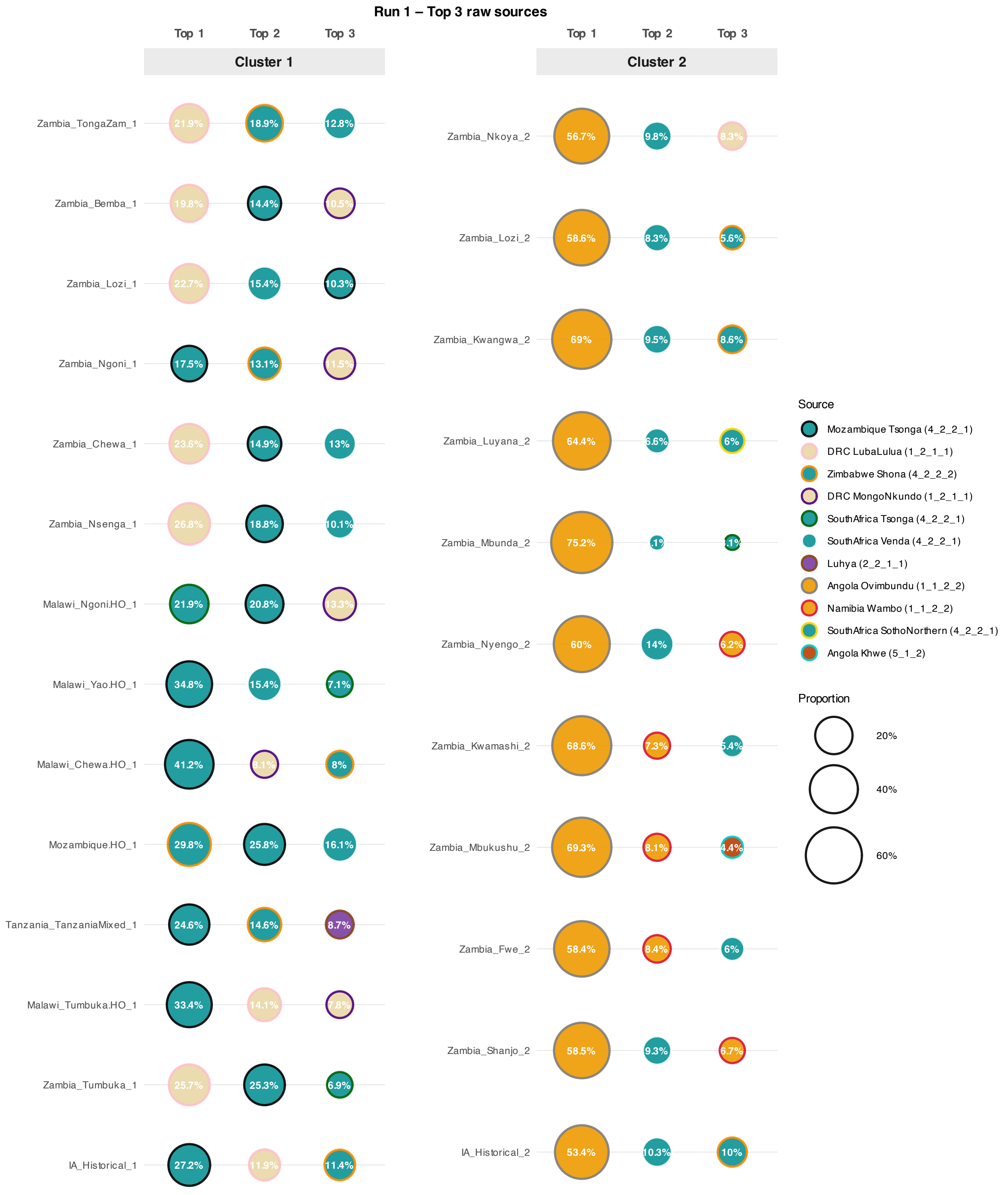

### Supplementary Figures 7

#### Run 2 – Top 3 raw sources

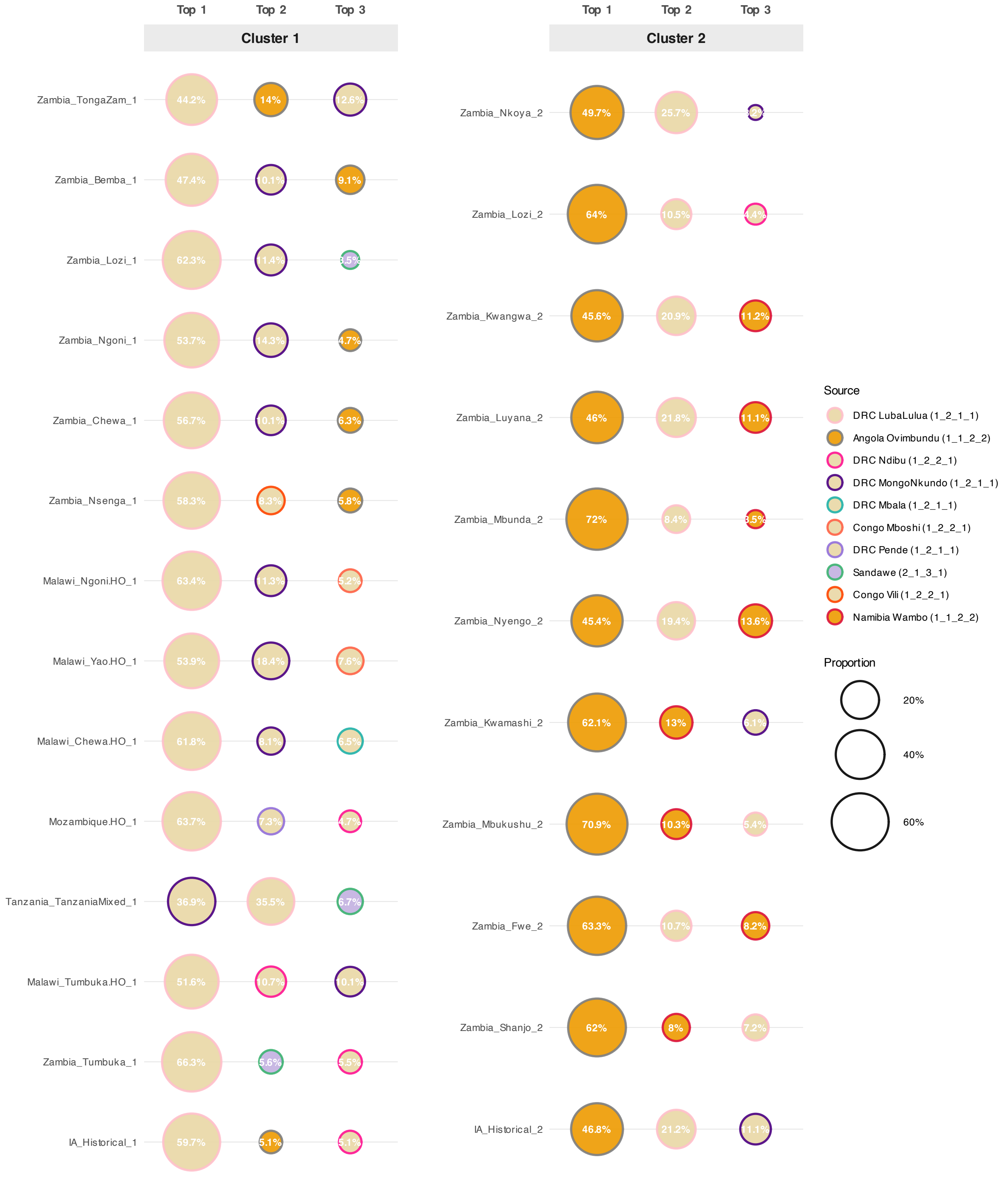

### Supplementary Figures 8

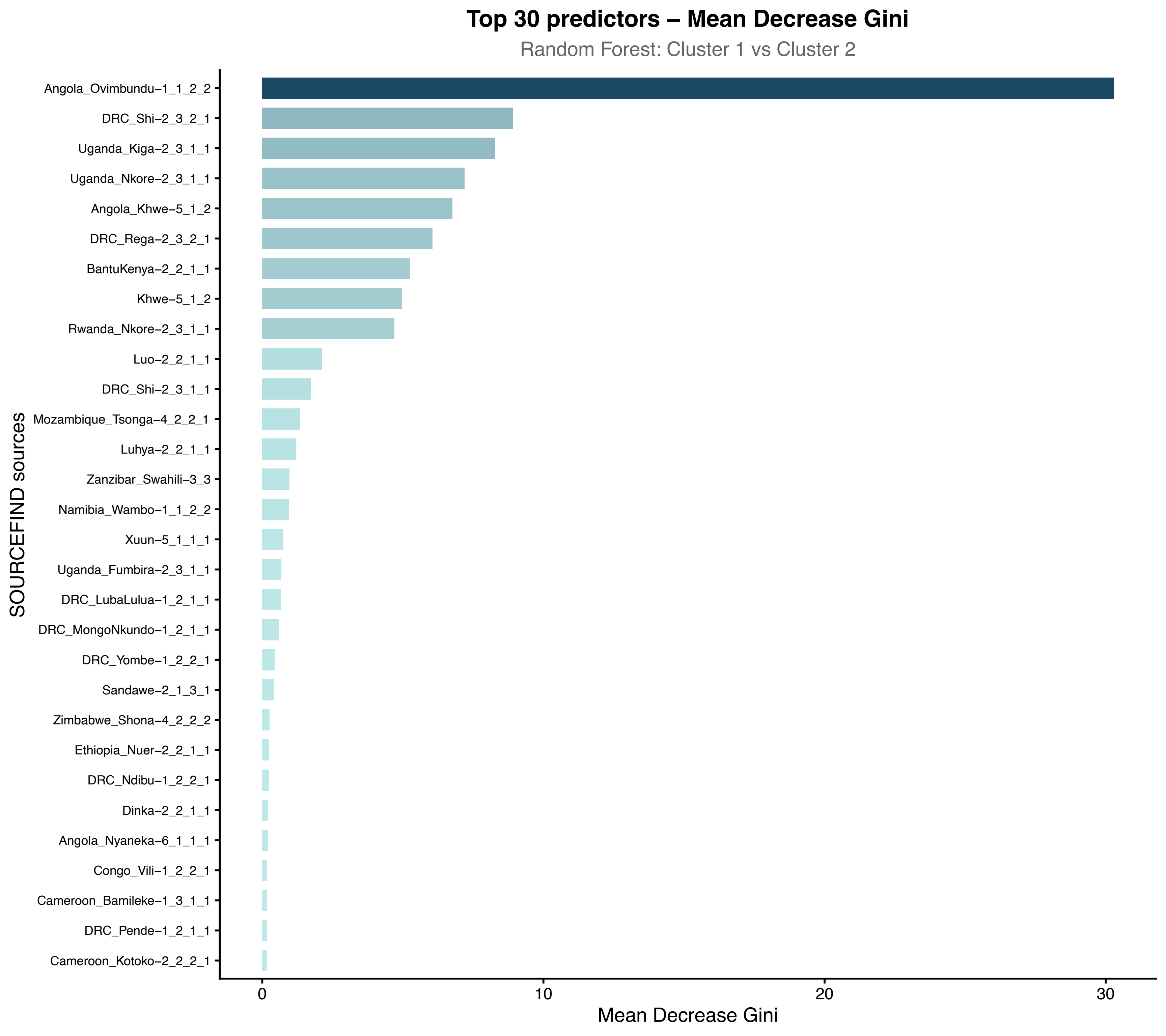

### Supplementary Figures 9

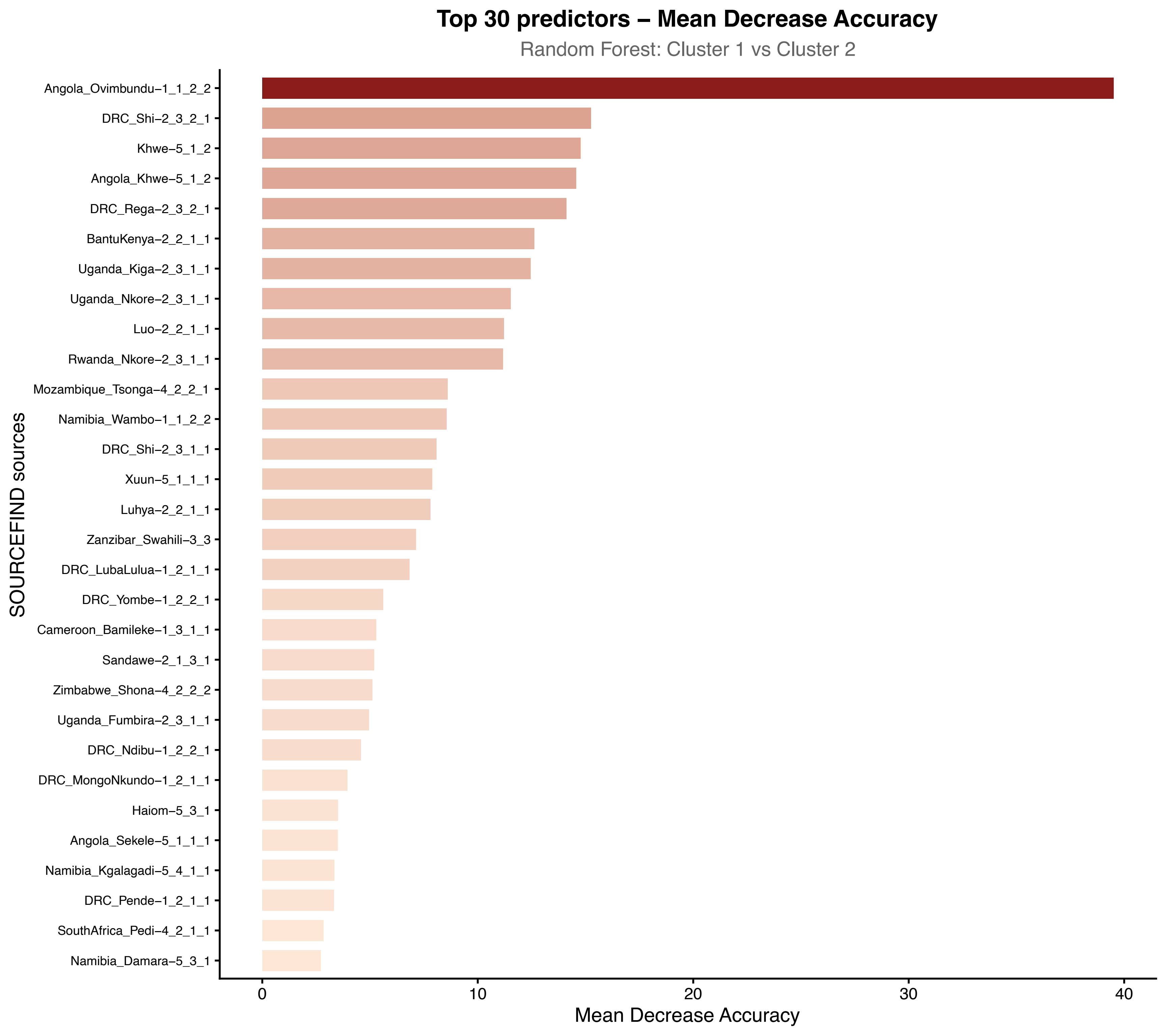

### Supplementary Figures 10

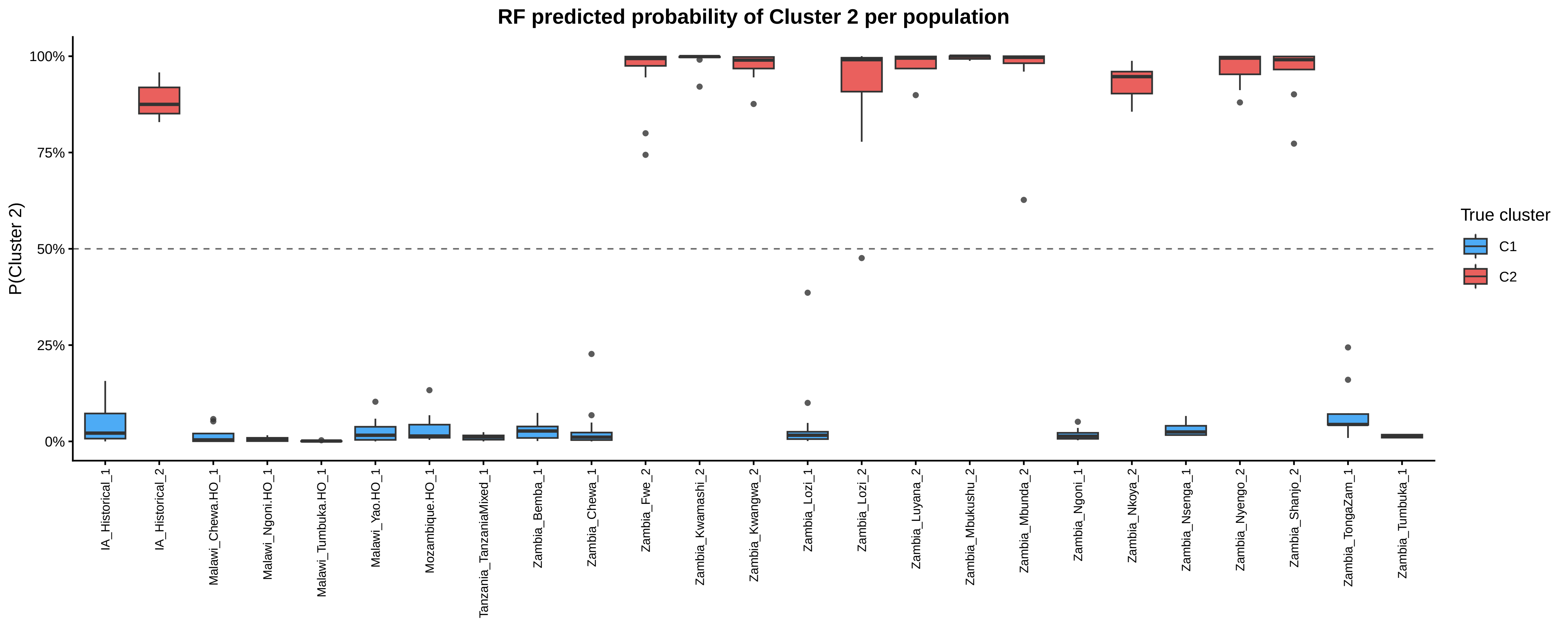

### Supplementary Figures 11

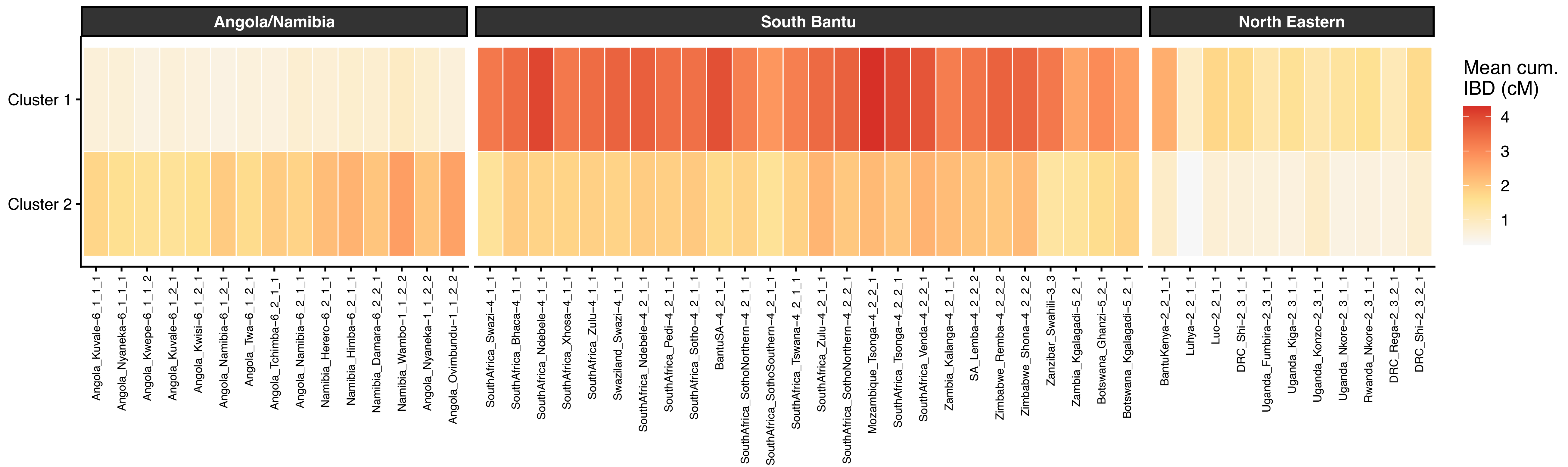
