## Supplementary Figures for "The south Congo Basin was critical to Bantu settlement of south central Africa"

**Supplementary Figure1.** PCA based on the chunck count matrix of Chromopainter (focused on modern south central individuals).

**Supplementary Figure 2.** Mean individual concordance across different number of clusters (K) identified by FineSTRUCTURE (FS_south central_). Concordance was estimated by comparing the state with the highest posterior probability (final state) with 100 MCMC samples.

**Supplementary Figure 3.** Proportion of individuals with membership concordance below 0.9 across different numbers of clusters (K) identified by FineSTRUCTURE (FS_south central_).

**Supplementary Figure 4.** TVD tree inferred from the ChromoPainter chunk length matrix using the algorithm described in the Methods section. After each merge, permutation tests were performed to assess genetic differentiation between the two resulting groups. Groups were considered statistically indistinguishable at p > 0.01.

**Supplementary Figure 5.** TVD matrix using the unlinked mode (-u switch) of Chromopainter.

**Supplementary Figure 6.** Top 3 source populations for each target group as inferred by SOURCEFIND (run1).

**Supplementary Figure 7.** Top 3 source populations for each target group as inferred by SOURCEFIND (run 2).

**Supplementary Figure 8. Top 30 SOURCEFIND donor populations ranked by Mean Decrease Gini.** Bar chart showing the 30 most important predictor variables in the Random Forest model discriminating Cluster 1 from Cluster 2, ranked by Mean Decrease Gini. Higher values indicate greater contribution to node purity across all decision trees. Colors reflect the magnitude of importance (light blue = low, dark blue = high).

**Supplementary Figure 9. Top 30 SOURCEFIND donor populations ranked by Mean Decrease Accuracy.** Bar chart showing the 30 most important predictor variables in the Random Forest model discriminating Cluster 1 from Cluster 2, ranked by Mean Decrease Accuracy. Higher values indicate a greater decrease in classification accuracy when the variable is permuted, reflecting its contribution to predictive performance. Colors reflect the magnitude of importance (light red = low, dark red = high).

**Supplementary Figure 10. Random Forest predicted probability of Cluster 2 membership per population.** Boxplots showing the distribution of individual-level predicted probabilities of belonging to Cluster 2 across all groups. Each box represents the interquartile range, with the median line and individual outliers shown. Colours indicate the true cluster assignment: blue (Cluster 1) and red (Cluster 2).

**Supplementary Figure 11. Differential IBD sharing between south central African clusters and modern Bantu-speaking populations across sub-Saharan Africa.** Mean cumulative IBD sharing (≥4 cM) between Cluster 1 and Cluster 2 and modern Bantu-speaking populations from three African geographic regions Each cell represents the mean IBD averaged across all possible individual pairs between a given cluster and an external population, including pairs sharing zero IBD. Cluster 1 shows markedly higher IBD with South Bantu and North Eastern populations, while Cluster 2 shows comparatively higher IBD with Angola/Namibia populations, reflecting distinct geographic affinities of the two clusters.
